# Fetal Thymic Expression Defines the Immunogenicity of Tumor Associated Antigens

**DOI:** 10.64898/2026.01.25.701584

**Authors:** Adrienne H. Long, Kaitlyn R. Tooker, Nicholas M. Pathoulas, Casey Beppler, Tatjana Bilich, Vishnu Shankar, Nathan A. Bracey, Mark M. Davis

**Affiliations:** Division of Hematology, Oncology, Stem Cell Transplantation and Regenerative Medicine, Department of Pediatrics, Stanford University School of Medicine, Stanford, CA, USA; Institute for Immunity, Transplantation and Infection, Stanford University School of Medicine, Stanford, CA, USA; Howard Hughes Medical Institute, Stanford University, Stanford, CA, USA; Department of Microbiology and Immunology, Stanford University School of Medicine, Stanford, CA, USA

**Keywords:** tumor associated antigens, cancer immunotherapy, immune checkpoint blockade, thymic tolerance

## Abstract

Tumor associated antigens (TAAs) are promising targets for cancer immunotherapy, yet their immunogenicity varies widely and remains poorly understood. Here, we show that the immunogenic potential of TAAs is largely shaped by their expression in the thymus, likely established during fetal development. By integrating single-cell transcriptomic data across fetal and postnatal thymic epithelial cells (TECs), we find that TAA expression in fetal TECs predicts both *in vitro* CD8⁺ T cell immunogenicity and the immune pressure against TAAs that occurs clinically in patients treated with immune checkpoint inhibitors. Notably, CD8⁺ T cells recognizing TAAs highly expressed in the fetal thymus exhibit attenuated transcriptional, signaling, and metabolic responses even in the naïve state, consistent with a tolerogenic imprinting imparted during thymic development and selection. Further, fetal thymic expression of TAAs can be leveraged to prioritize candidate targets for therapeutic use. These findings provide a biologic basis for the extreme variability seen in TAA immunogenicity and define guiding principles for rational antigen selection to drive the next generation of effective TAA-targeted immunotherapies.

## INTRODUCTION

Novel immunotherapies are urgently needed for cancers where survival remains poor despite optimized chemotherapy. Recent work has focused largely on neoantigens, mutated cancer-specific proteins recognized as foreign by the immune system.^1–3^ However, these targets are infrequent in tumors with lower mutational burdens, such as hematologic and pediatric malignancies, and are often unique to individual patients.^4,5^ Tumor associated antigens (TAAs) have long represented an important, alternative class of potential targets. TAAs are non-mutated self-proteins that are aberrantly expressed in cancers compared to healthy postnatal tissues. TAAs are appealing targets because they are far more abundant than neoantigens within a given tumor and are often shared across different patients and cancer types.^6,7^ Although several TAA-targeted therapies have shown meaningful clinical benefit, most have produced limited responses.^8–11^ The reason why targeting some TAAs yields clinical success, while targeting others does not, remains poorly understood.

Currently, most TAA target selection focuses on identifying which TAAs have the highest expression on tumor compared to healthy tissues.^12^ Far less attention is given to whether TAAs are equally immunogenic from the perspective of the immune system. As non-mutated self-antigens, TAAs are likely subject to immune tolerance, but the extent to which this occurs remains poorly defined. T cell tolerance is first established in the thymus, where thymic epithelial cells (TECs) present a diverse array of self-antigens to developing thymocytes, eliminating strongly self-reactive clones and shaping the mature T cell repertoire.^13^ Historically, the vast majority of self-reactive CD8⁺ T cells were thought to be eliminated through this process, known as clonal deletion, which should severely limit immunologic responses to TAAs. However, we and others have demonstrated that a surprisingly large number of self-reactive CD8⁺ T cells evade thymic deletion and populate the periphery.^14,15^ This unexpected escape challenges the traditional view of thymic tolerance and raises fundamental questions about how CD8^+^ T cells reactive to TAAs are controlled. Understanding the regulation of CD8⁺ T cell responses to TAAs is critical to harnessing these cells for the development of effective TAA-directed immunotherapies.

Here, we demonstrate that TAAs exhibit striking variability in their inherent immunogenicity, which we trace to thymic tolerance established early in human development. By integrating single-cell transcriptomic profiles of fetal and postnatal TECs with functional assays of CD8⁺ T cell responses to TAAs, we show that the level and developmental timing of thymic TAA expression dictates their immunogenicity. TAAs with low fetal TEC expression disproportionately elicit robust CD8⁺ T cell responses, whereas those highly expressed during fetal thymic development are associated with tolerance and poor immune recognition.

Furthermore, this tolerance extends beyond clonal deletion, as CD8⁺ T cells specific for TAAs highly expressed in the fetal thymus are not completely deleted, but persist in a hypofunctional state driven by a restrained transcriptional program. These findings reveal that fetal thymic tolerance not only prunes the TAA-specific CD8⁺ T cell repertoire, but also durably programs the functional quality of the TAA-reactive response.

Building on this framework, we extend our findings to show that fetal thymic tolerance has direct implications for tumor immune recognition and immunotherapy design. Leveraging public datasets from patients treated with immune checkpoint inhibitors, we find that tolerance imparted by fetal thymic expression of TAAs shapes the immune pressures exerted on tumors during therapy. Further, incorporating fetal thymic expression patterns into antigen selection strategies enhances the ability to select TAAs that are most likely to escape tolerance and elicit potent antitumor responses. Together, our findings define a developmental foundation for TAA immunogenicity and establish an important new framework for identifying and targeting the most immunogenic self-antigens for cancer immunotherapy.

## RESULTS

### Immunogenicity of commonly targeted TAAs is highly variable

As an initial step towards understanding the immunogenicity of TAAs, we compared the CD8^+^ T cell responses to a panel of commonly targeted HLA-A*02-restricted TAAs, all of which have been extensively studied in clinical trials (Extended Data Fig 1A).^16–20^ Nominally healthy PBMCs were stimulated with TAA peptides, and antigen-specific CD8^+^ T cell responses quantified using peptide-MHC tetramers (Fig 1A).^21^ We found that a subset of TAAs, including MART-1 and MAGE-A3, consistently drove robust expansion of antigen-specific CD8⁺ T cells (Fig 1B). In contrast, other TAAs such as PMEL, WT1, and PRAME elicited only modest responses. These differences could not be fully attributed to baseline differences in unstimulated T cell frequencies, as we also found MART-1 and MAGE-A3 peptides drove higher rates of proliferation than the other TAAs (Fig 1C), suggesting instead an intrinsic difference in TAA reactivity. To confirm this finding, we compared other functional aspects of CD8⁺ T cell responses to the TAA panel. Consistent with the proliferation data, highly immunogenic TAAs such as MART-1 and MAGE-A3 induced strong upregulation of activation markers CD38 and CD25 (Fig 1D-E) and robust degranulation (Fig 1F) with peptide stimulation, whereas other TAAs triggered only limited responses. Taken together, these findings demonstrate that while TAAs are all self-antigens, they differ markedly in their capacity to elicit strong CD8⁺ T cell responses.

**Fig 1:**
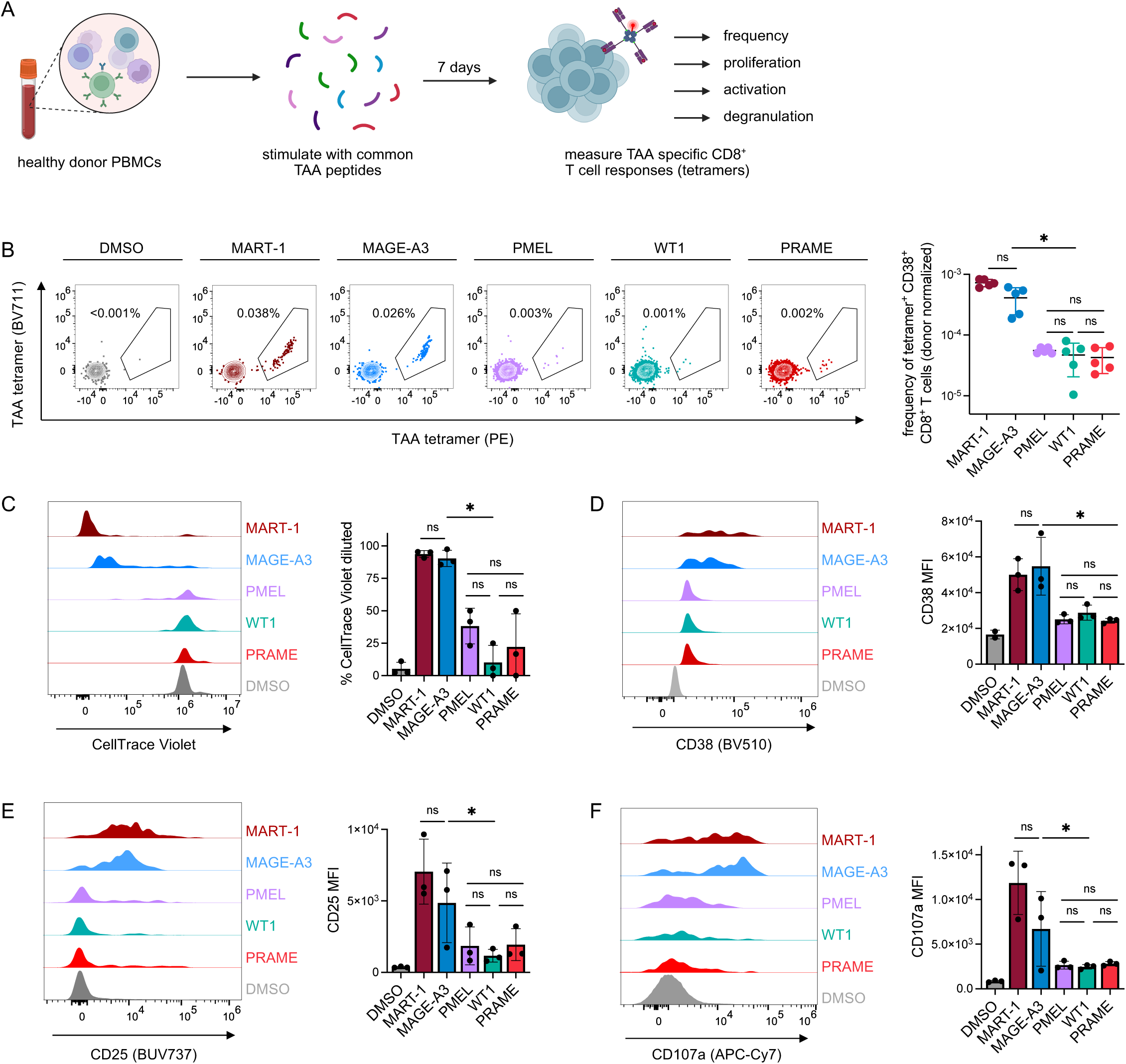
Immunogenicity of TAAs is highly variable. **(A)** Experimental design for evaluating CD8^+^ T cell responses of TAAs. Healthy donor PBMCs were stimulated with TAA peptides, cultured for 7 days, and antigen-specific responses measured with peptide-MHC tetramers using flow cytometry. **(B)** Frequency of activated TAA-specific CD8^+^ T cells (tetramer^+^ CD38^+^) after peptide stimulation. Left, representative staining; right, quantification across n = 5 donors, showing that some TAAs elicit robust responses whereas others fail to expand. **(C)** Proliferation of TAA-specific CD8⁺ T cells after peptide stimulation, measured by dilution of CellTrace Violet staining, indicating that differences in proliferation after peptide stimulation contribute to heterogeneity in TAA responses. **(D)** Expression of the activation marker CD38 on activated TAA-specific CD8⁺ T cells after peptide stimulation, demonstrating that strongly immunogenic TAAs drive higher activation levels. **(E)** Expression of the activation marker CD25 on activated TAA-specific CD8⁺ T cells after peptide stimulation, demonstrating that strongly immunogenic TAAs drive higher activation levels. **(F)** Degranulation of activated TAA-specific CD8⁺ T cells after peptide stimulation, measured by CD107a externalization, demonstrating those TAAs that induce robust responses also induce greater degranulation after peptide stimulation. For panels (c-f), left plots show representative FACS traces and right plots show quantification across n = 3 donors. A total of 12 unique donors were analyzed across all panels. * represents p < 0.05, as measured by paired t-test.

### Immunogenicity of TAAs is regulated by thymic tolerance established very early in life

By definition, TAAs have limited expression in healthy tissue. Therefore, we hypothesized that thymic tolerance, rather than peripheral tolerance, might be key to regulating TAA immunogenicity. Thymic tolerance is established during thymopoiesis through presentation of self-antigens by thymic epithelial cells (TECs; Fig 2A).^13^ Thus, to assess whether this form of tolerance contributes to TAA immunogenicity, we analyzed a single-cell thymus atlas^22^ and quantified TAA expression across TECs. We analyzed 17,480 TECs from 16 donors spanning fetal to adult life (10 weeks post-conception to 35 years; Fig 2B and Extended Data Fig 2A-B). Using an expanded panel of TAAs and other self-antigens with restricted healthy tissue expression (Extended Data Fig 1A), we compared each antigen’s TEC expression with its immunogenicity, defined as the frequency of tetramer-positive CD8⁺ T cells after peptide stimulation (Extended Data Fig 1B-D). Because reduced thymic exposure should lead to weaker tolerance, we hypothesized that TAAs with lower TEC expression would exhibit greater immunogenicity.

**Fig 2:**
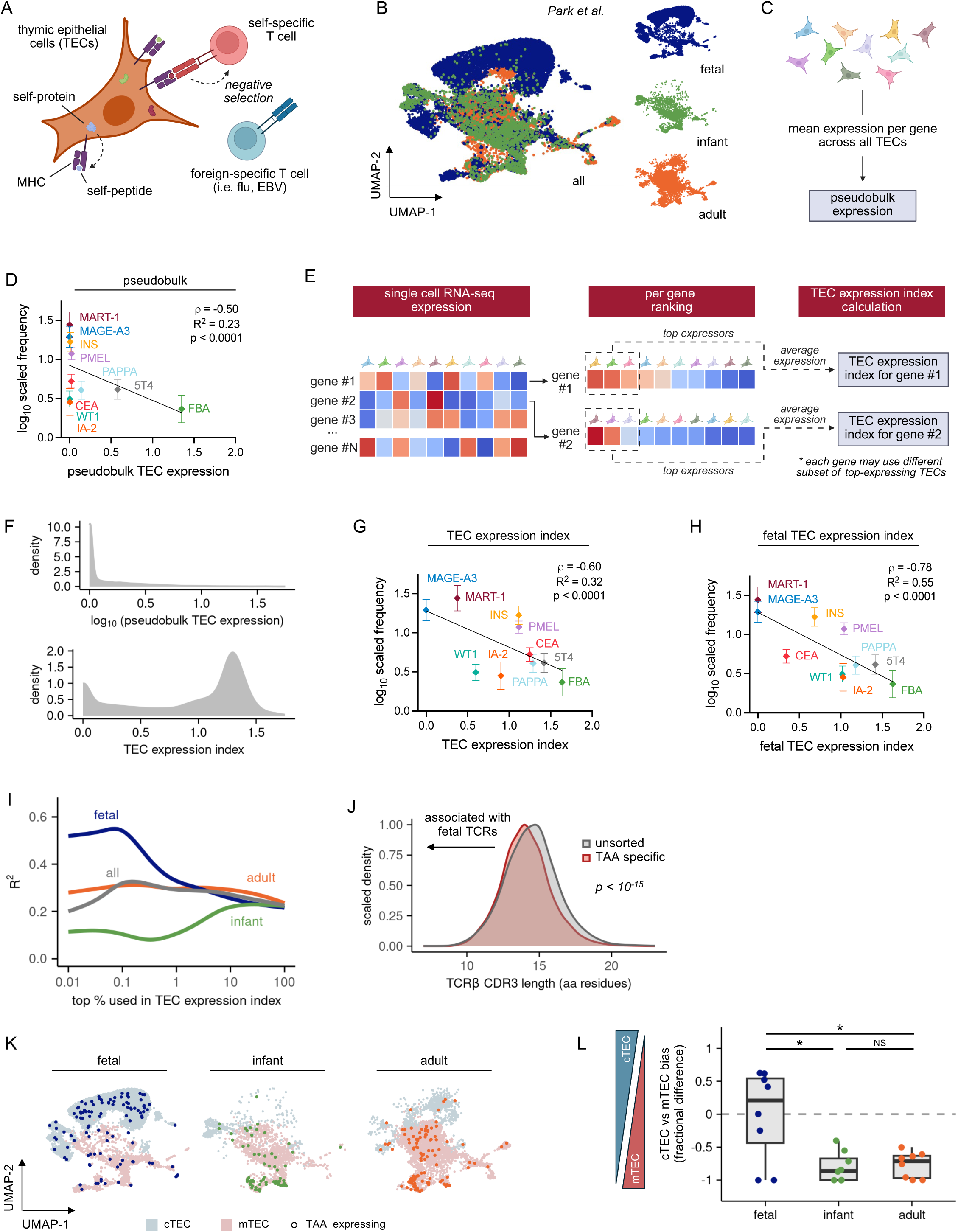
Immunogenicity of TAAs is regulated by thymic tolerance established in early life. **(A)** Schematic of thymic tolerance mediated by thymic epithelial cells (TECs). Thymic tolerance is established through TEC expression of self-antigens, enabled by promiscuous gene expression of non-thymic, tissue-restricted proteins. This stochastic expression allows TECs to present diverse self-antigens on MHC molecules, bringing potentially autoreactive T cells into contact with their cognate antigens and triggering negative selection. **(B)** UMAP representation of TEC transcriptomes, highlighting developmental stages within the data set: fetal (blue), infant (green), and adult (orange). Plots illustrate the marked shift in TEC composition and gene expression across ontogeny. **(C)** Schematic for pseudobulk analysis approach for TAA expression across all TECs. **(D)** Correlation between pseudobulk TAA expression across TECs (log-normalized expression values) and the frequency of TAA-specific CD8⁺ T cells, demonstrating that pseudobulk TEC expression has a moderate correlation with immunogenicity (ρ = –0.50) but overall limited predictive power. **(E)** Schematic for TEC expression index calculation. Single-cell RNAseq data from TECs were used to rank cells by expression for each gene independently. For every gene, the average log-normalized expression among the top-expressing TECs was then calculated, generating a gene-specific TEC expression index which reflects the expression within the subset of cells most actively transcribing the given gene. **(F)** Distribution of TEC expression for all transcribed genes in the human genome. Top, distribution of pseudobulk expression for all genes; bottom, distribution of TEC expression index values for all genes, highlighting that TEC expression index captures rare, highly expressed transcripts that are masked in pseudobulk averaging. **(G)** Correlation between TEC expression index (top 0.2% expressors across all TECs) and the frequency of TAA-specific CD8⁺ T cells, demonstrating that the TEC expression index better predicts TAA immunogenicity than pseudobulk TAA expression. **(H)** Correlation between fetal TEC expression index (top 0.06% expressors within fetal TECs) and the frequency of TAA-specific CD8⁺ T cells, demonstrating that fetal TEC expression provides the strongest predictive signal for TAA immunogenicity. **(I)** Impact of TEC ontogeny on the ability of TEC expression index to predict TAA immunogenicity. TEC single-cell data were first stratified into fetal, infant, adult, or all TECs, and TEC expression indices calculated for the top 0.01-100% expressors (100% = pseudobulk). The fetal TEC expression index calculated from 0.06% of the top expressors provides the strongest predictive signal for TAA immunogenicity. P.adj (Bonferroni correction) for all points are < 0.05, except infant top % expressors between 0.1-1. **(J)** Comparison of TCRβ CDR3 length distributions between TAA-specific CD8⁺ T cells versus unsorted CD8⁺ T cells from the same donors. CDR3 lengths were calculated from sequences obtained using oligo-tagged tetramers with single-cell RNAseq. Results reveal that TAA-specific TCR repertoires are enriched for shorter CDR3s, consistent with fetal origin. Statistically compared with Wilcoxon Rank Sum test. **(K)** UMAP visualization showing the distribution of TAA-expressing TECs across cTEC (light blue) and mTEC (light red) compartments, for fetal, infant, and adult TEC populations, demonstrating a skewing of TAA-expressing fetal TECs towards the cTEC subtype, and skewing towards mTECs for infant and adult TECs. For each TAA, up to 50 expressing TECs were randomly sampled and overlaid as individual dots. **(L)** Quantification of cTEC versus mTEC bias among TAA-expressing TECs across fetal, infant, and adult thymic populations, shown as fractional difference ([# cTEC – # mTEC]/[# cTEC + # mTEC]). Aggregated results show that fetal TECs exhibit a stronger cTEC bias compared to infant and adult TECs. Each point represents an individual TAA from the analyzed panel. TAAs not detected in TECs were excluded. All analyses in this figure were conducted on n = 12 donors. For correlation plots (d), (g) and (h), frequency reported is of TAA-specific CD8⁺ T cells measured by oligo-tagged tetramers with CITE-seq, normalized and scaled across donors; ρ denotes Spearman correlation while R^2^ and p denote linear regression results.

Quantifying TEC expression using a pseudobulk approach revealed a moderate inverse association between TAA expression in TECs and TAA immunogenicity (Fig 2C-D; Spearman ρ = –0.50; R² = 0.23), consistent with a role for thymic tolerance in shaping these responses. However, this approach provided limited ability to distinguish strongly from weakly immunogenic antigens, suggesting that averaging expression across all TECs may obscure the TEC subsets most relevant for tolerance.

Self-antigen expression in the thymus is highly restricted, with each gene being expressed by only a small fraction of TECs (<1%) at any given time.^23^ Thus, we reasoned that focusing on the highest-expressing TECs might better capture the tolerogenic signal. We therefore developed a “TEC expression index” that measures expression specifically in the top-expressing TECs for each gene, reflecting the cells most likely to mediate tolerance (Fig 2E). This biologically motivated metric more effectively captured meaningful variation in TEC expression. Whereas pseudobulk TEC expression values calculated across all genes in the genome produced a compressed, low-expression distribution, the TEC expression index revealed clear separation between genes with high versus low TEC expression (Fig 2F). When applied to our TAA panel, the TEC expression index preserved this improved separation, distinguishing antigens that were obscured by pseudobulk values and yielding a stronger correlation with immunogenicity (Fig 2G).

We next asked whether the developmental stage of the thymus influenced the association between TEC expression and TAA immunogenicity. Thymic tolerance undergoes marked transitions over fetal, infant, and adult life, with fetal thymopoiesis characterized by relatively permissive tolerance and immature expression of key tolerance pathways.^24^ Unexpectedly, stratifying TECs by developmental stage revealed that fetal TEC expression was the most predictive of TAA immunogenicity, explaining more than half of the variance (R² = 0.55; Fig 2H–I). This relationship was robust to differences in sequencing depth, cell number, and approach to quantifying TAA immunogenicity (Extended Data Fig 1E-G). Taken together, these results demonstrate that TEC expression patterns predict TAA immunogenicity, with fetal TEC expression providing the greatest predictive strength among developmental stages.

The correlation between TAA immunogenicity and expression in the fetal thymus suggested that TAA-specific CD8^+^ T cells were educated by fetal TECs and thus also fetally derived. We therefore asked whether TAA-specific CD8^+^ T cells carry intrinsic features consistent with a fetal origin. Fetal T cells are known to have T cell receptors (TCRs) with shortened CDR3 regions, the most hypervariable portion of the receptor, because expression of terminal deoxynucleotidyl transferase (TdT) is delayed until late in gestation (after approximately 20 weeks).^25^ Using oligo-tagged tetramers and single-cell TCR sequencing, we found that TAA-specific CD8^+^ T cells consistently exhibited shorter TCRβ CDR3 regions compared to bulk CD8⁺ T cells from the same donors (Fig 2J; Extended Data Fig 1H). These results provide independent evidence that TAA-specific T cells may arise during fetal thymopoiesis.

This finding prompted us to ask whether distinctive features of fetal TECs might underlie their enhanced ability to predict TAA immunogenicity. A defining characteristic of fetal thymus is its distinct TEC composition, which shifts markedly over development (Extended Data Fig 2C-D).^22,26^ In the postnatal thymus, cortical TECs (cTECs) guide early T cell maturation by ensuring ability to bind to MHC, whereas medullary TECs (mTECs) eliminate strongly self-reactive cells to enforce tolerance, a process driven largely by the transcription factor AIRE. In contrast, the fetal thymus is dominated by cTECs, with only rare AIRE^low^ mTECs (Extended Data Fig. 2E).^22,26^ Surprisingly, TAA expression in the fetal thymus was concentrated in cTECs rather than mTECs, whereas postnatal TAA expression was almost entirely mTEC-restricted (Fig 2K-L). This suggests that fetal cTECs can provide tolerogenic cues even before the mature medullary tolerance machinery develops. Notably, restricting analyses to cTECs within the infant or adult thymus did not restore the predictive strength observed in fetal TECs (Extended Data Fig 2F), indicating that the superior tolerogenic capacity of fetal TECs reflects a distinct developmental state rather than cTEC abundance alone. Together, these data suggest a previously unrecognized fetal-specific thymic program in which cTECs play an unexpectedly prominent role in establishing tolerance within TAA-specific CD8^+^ T cells.

### Fetal TEC expression predicts immunogenic TAA dropout with checkpoint inhibitor therapy

Our initial findings suggested that fetal TEC expression can be used to predict TAA immunogenicity, but these studies were limited to healthy donor PBMCs stimulated with peptides. To independently validate this observation in a cancer-specific context, we turned to published datasets from melanoma patients treated with checkpoint inhibitors.^27–29^ Recent work has proposed that immune recognition of non-mutated cancer antigens, in addition to neoantigens, contributes to clinical responses to checkpoint blockade.^7^ This was demonstrated through analyses showing that expression of non-mutated antigens “drops out’’ in tumor biopsies from patients responding to treatment, consistent with immune-mediated elimination of antigen-expressing tumor cells. However, that work primarily focused on cancer antigens aberrantly translated from non-coding regions and evaluated only antigens not expressed by TECs. As a result, it provided limited insight into how thymic tolerance to conventional, coding-region TAAs might also shape responses to checkpoint blockade.

To determine whether conventional, coding-region TAAs are similarly subject to immune pressure during checkpoint blockade, we analyzed three RNAseq datasets from paired biopsies obtained from melanoma patients collected before and during checkpoint inhibitor therapy (Riaz et al., n = 43; Gide et al., n = 16; Du et al., n = 4).^27–29^ For each patient at both timepoints, we identified TAA candidates and predicted epitopes presented by class I HLA molecules (Fig 3A). We then compared TAA epitope abundance pre– and on-therapy within each patient to identify dropout events indicative of immune pressure. Across studies, responders lost an average of 20 ±10% of predicted TAA epitopes following therapy (95% CI; p < 0.0001), whereas non-responders showed no significant loss of epitopes (−3 ± 7%, 95% CI; Fig 3B-C). These findings suggest that immune pressure against traditional TAAs, in addition to what has been previously shown with antigens arising from non-coding regions, contributes to responses to checkpoint inhibitor therapy.

**Fig 3:**
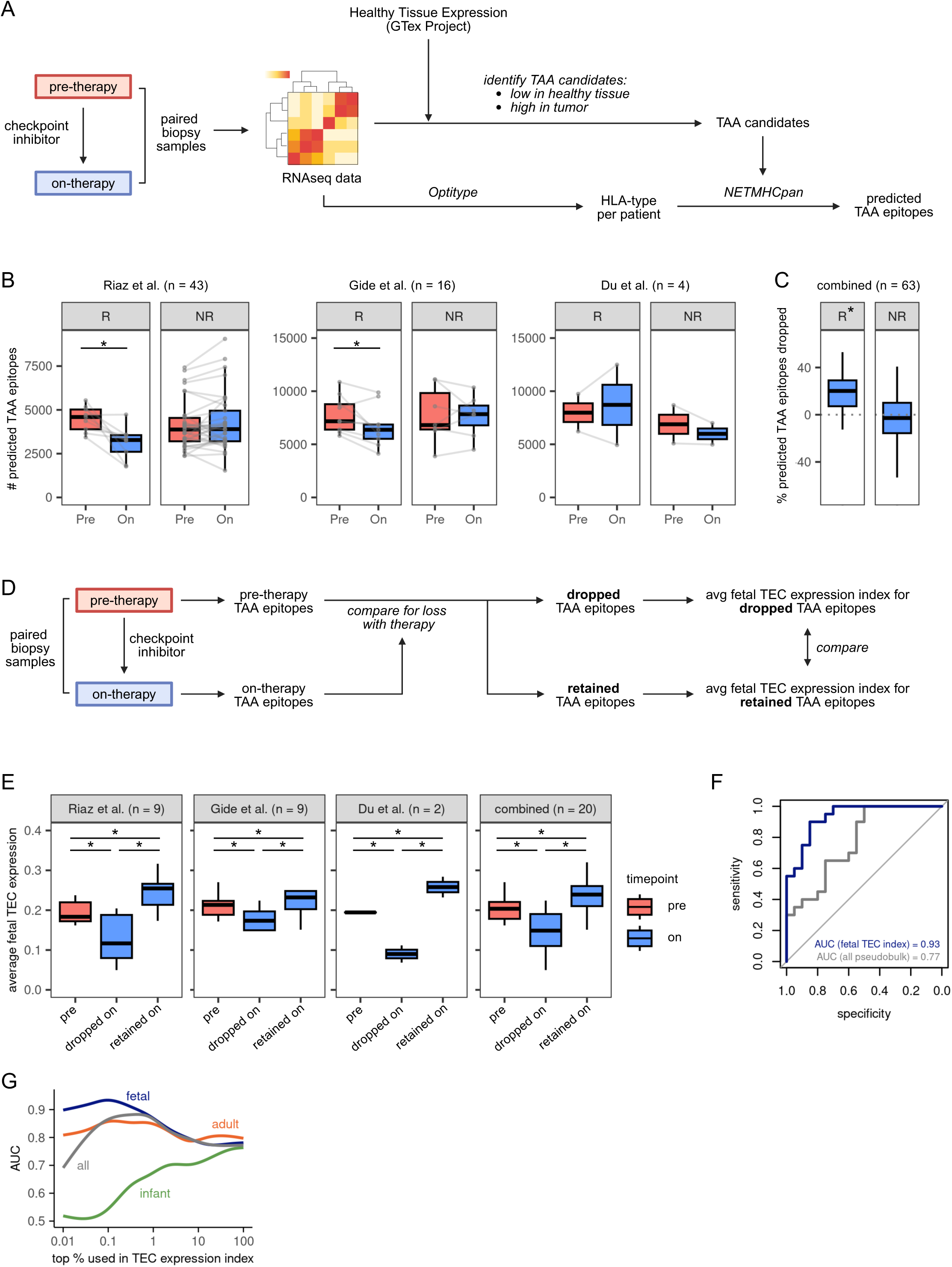
Fetal TEC expression predicts immunogenic TAA dropout with checkpoint inhibitor therapy. **(A)** Experimental design for evaluating TAA dropout during checkpoint inhibitor therapy. Paired pre– and on-treatment melanoma RNAseq datasets were analyzed to identify TAAs with high tumor and low healthy tissue expression (GTEx reference). HLA types were inferred using OptiType, and TAA epitopes predicted to be presented by patient-specific HLAs were identified with NetMHCpan. Predicted TAA epitopes were then compared between pre– and on-therapy samples to identify antigens potentially lost under immune pressure. **(B)** Number of predicted TAA epitopes presented in patient tumors before and during checkpoint inhibitor therapy. Data are stratified by clinical response (responders, R, vs. non-responders, NR), revealing greater epitope loss among responders. Data shown for three independent melanoma datasets. **(C)** Percentage of predicted TAA epitopes that were dropped with checkpoint inhibitor therapy across all three melanoma datasets, demonstrating consistent antigen loss with effective therapy. **(D)** Schematic approach for evaluating fetal TEC expression of TAAs that are differentially dropped versus retained during checkpoint inhibitor therapy. Predicted TAA epitopes were identified from paired pre– and on-therapy tumor RNAseq samples. Epitopes present pre-therapy but absent on-therapy were classified as dropped, while those present in both were classified as retained. Fetal TEC expression indices were then calculated for each TAA, and compared between dropped and retained epitope groups. **(E)** Average fetal TEC expression of pretreatment TAA epitopes per patient, stratified by whether antigens identified were subsequently dropped or retained on treatment. Analyses are shown for three independent melanoma datasets and for all datasets combined, demonstrating that dropped antigens consistently exhibit lower fetal TEC expression than retained antigens. **(F)** Receiver operator characteristic (ROC) curve for using average TEC expression indices to separate dropped versus retained predicted TAA epitopes with checkpoint inhibitor therapy, demonstrating superior predictive performance of the fetal TEC expression index. Blue, analysis performed with fetal TEC expression index from top 0.06%; grey, analysis performed with pseudobulk TEC expression from all TECs. **(G)** Impact of TEC ontogeny on the ability of TEC expression index to distinguish dropped versus retained predicted TAA epitopes with checkpoint inhibitor therapy. TEC single-cell data were first stratified into fetal, infant, adult, or all TECs, and TEC expression indices calculated for the top 0.01-100% expressors (100% = pseudobulk). Area under the curve (AUC) reported represents outcome of ROC analyses for ability of average TEC expression to stratify dropped versus retained TAA epitopes with checkpoint inhibitor therapy. Results demonstrate that the fetal TEC expression index provides the greatest ability to separate the average TEC expression index between dropped and retained TAAs per patient. All * represent p <0.05 measured by paired Wilcoxon Rank Sum test.

We next asked whether fetal TEC expression of TAAs influences their likelihood of dropout during checkpoint inhibitor therapy. Predicted TAA epitopes present in pretreatment samples were classified as either lost (dropped out) or retained following therapy, and the fetal TEC expression index was calculated for each group (Fig 3D). Across all three studies, TAAs that dropped out with clinical response to checkpoint inhibitors had significantly lower fetal TEC expression indices compared with retained TAAs (Fig 3E), indicating that thymic tolerance to traditional TAAs shapes the immune pressure applied to distinct TAAs during checkpoint inhibitor therapy. Notably, consistent with our peptide-stimulation assays in healthy donor PBMCs, average fetal TEC expression provided the greatest ability to distinguish dropped from retained TAAs (Fig 3F-G), supporting the conclusion that thymic tolerance established early in life plays a critical role in regulating clinically relevant immune responses to TAAs.

### Thymic imprinting constrains activation of TAA-specific CD8⁺ T cells

Thymic tolerance is traditionally viewed through the lens of clonal deletion, though more recent studies in both humans and mice have demonstrated that many self-reactive T cells escape negative selection yet remain functionally restrained.^14,30,31^ Importantly, these studies analyzed T cells with mixed naïve and memory phenotypes, making it difficult to determine whether their reduced responsiveness arose from thymic imprinting or peripheral tolerance. Our finding that TAA immunogenicity correlates with fetal thymic expression suggested that thymic experience may play a critical role by imprinting a lasting functional difference on surviving CD8^+^ T cells specific for TAAs. To test this, we performed multimodal single-cell profiling of TAA-specific CD8⁺ T cells using oligo-tagged tetramers (extended TAA panel), both when unstimulated and following peptide stimulation for 7 days (Fig 4A, Extended Data Fig 1A, Extended Data Fig 3). This single-cell approach enabled resolution of individual tetramer specificities, allowing analysis of rare antigen-specific populations that would otherwise require pooling for bulk analyses and thus loss of specificity-resolved information. We focused on the naïve compartment (Fig 4B, Extended Data Fig 4A; defined by overlying CD45RA^+^ CD45RO^−^ protein expression onto unsupervised RNA-level clustering), to isolate the contribution of thymic tolerance and minimize potential effects from peripheral antigen exposure.

**Fig 4:**
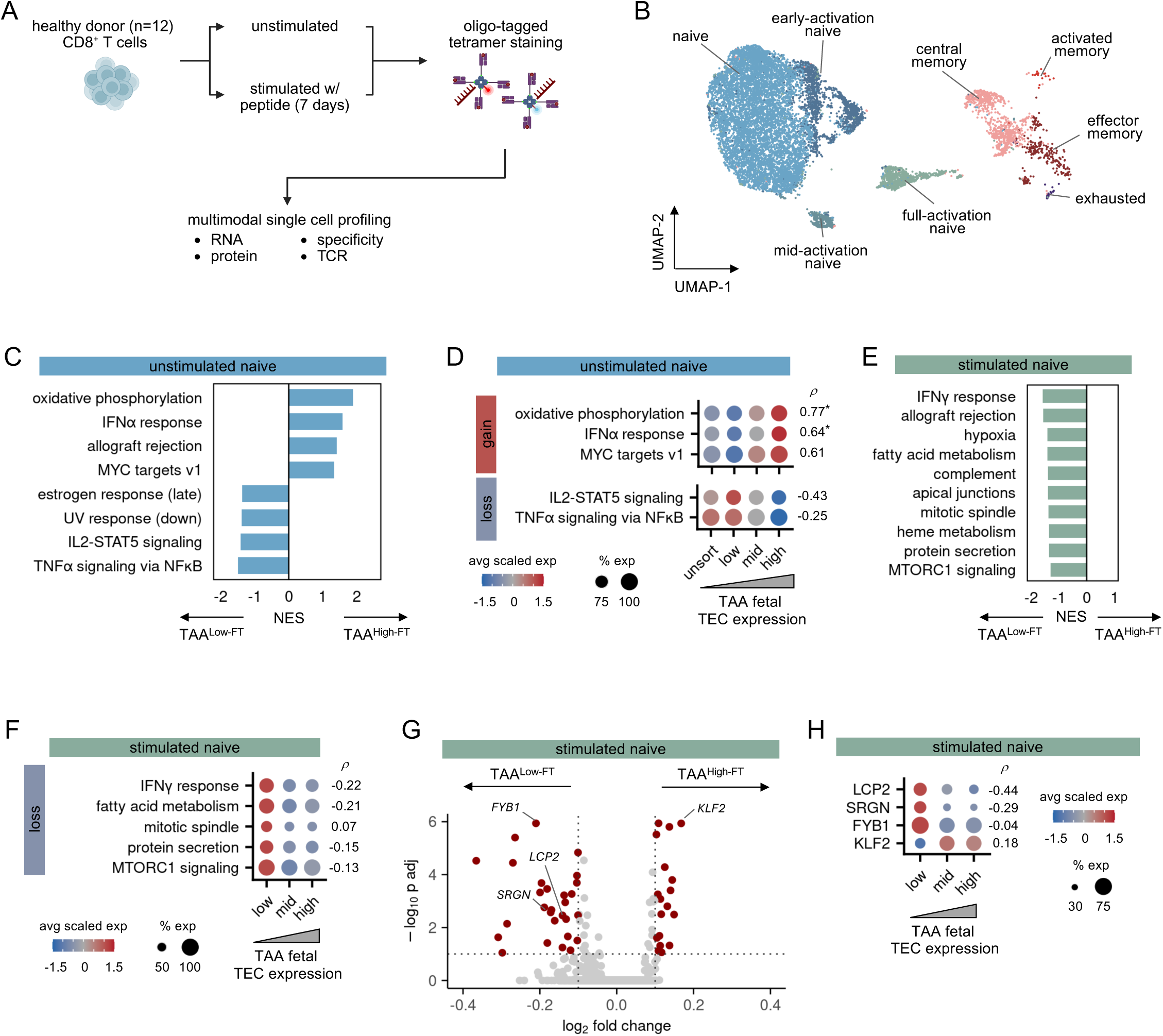
Thymic imprinting constrains activation of TAA-specific CD8⁺ T cells. **(A)** Schematic of single cell evaluation of TAA-specific CD8⁺ T cells. Peripheral blood T cells from healthy donors were labeled with oligo-tagged tetramers to define antigen specificity, flow-sorted, and subjected to multimodal single-cell analysis, including RNA sequencing, surface protein profiling (CITE-seq), and paired TCR sequencing. Matched unstimulated and 7-day peptide-stimulated samples were analyzed to characterize transcriptional and phenotypic states of TAA-specific CD8⁺ T cells. **(B)** UMAP of single-cell transcriptomes from TAA-specific CD8⁺ T cells. Integrated data from unstimulated and stimulated samples are shown. Cells are colored by memory phenotype, revealing that most TAA-specific cells reside within a naïve-like state, indicating limited prior activation. **(C)** GSEA analysis of unstimulated, TAA-specific naïve CD8⁺ T cells comparing those targeting TAA^High-FT^ versus TAA^Low-FT^ (fetal TEC expression indices > 0.75 or < 0.75, respectively; FDR <0.1). Naïve CD8^+^ T cells targeting TAA^High-FT^ have gained and lost activity in key T cell functionality pathways. **(D)** Module score analysis of unstimulated, TAA-specific naïve CD8⁺ T cells. Pathway module scores derived from GSEA leading-edge genes. T cells were binned into those targeting TAA^Low-FT^, TAA^Mid-FT^, and TAA^High-FT^ (fetal TEC expression <0.5, 0.5-1.1, and >1.1) to illustrate graded trends. Module expression correlated with fetal TEC expression index, demonstrating a relationship between thymic expression and pathway activity. **(E)** GSEA analysis of peptide-stimulated, TAA-specific naïve CD8⁺ T cells comparing those targeting TAA^High-FT^ versus TAA^Low-FT^ (fetal TEC expression indices > 0.75 or < 0.75, respectively; FDR <0.1). Naïve CD8^+^ T cells targeting TAA^Low-FT^ demonstrate a significantly higher enrichment for T cell activation pathways. **(F)** Module score analysis of peptide-stimulated, TAA-specific naïve CD8⁺ T cells. Pathway module scores derived from GSEA leading-edge genes. T cells were binned into those targeting TAA^Low-FT^, TAA^Mid-FT^, and TAA^High-FT^ (fetal TEC expression <0.5, 0.5-1.1, and >1.1) to illustrate graded trends. Activation signatures were preferentially induced in low fetal-TEC expressors, indicating a threshold-like relationship between thymic exposure and activation capacity. **(G)** Differential gene expression across peptide-stimulated, TAA-specific naïve CD8⁺ T cells. Volcano plot comparing CD8⁺ T cells targeting TAA^High-FT^ vs TAA^Low-FT^. Activation-associated genes were enriched in T cells targeting TAA^Low-FT^, whereas T cells targeting TAA^High-FT^ retained quiescence markers and showed reduced adaptor gene expression consistent with imprinted tolerance. **(H)** Expression of select genes differentially regulated between T cells targeting TAA^Low-FT^ versus TAA^High-FT^. T cells were binned into those targeting TAA^Low-FT^, TAA^Mid-FT^, and TAA^High-FT^ (fetal TEC expression <0.5, 0.5-1.1, and >1.1) to illustrate graded trends. T cells targeting TAA^High-FT^ showed higher expression of quiescence genes (KLF2) and lower expression of cytolytic genes and TCR signaling adaptors (SRGN, FYB1, LCP2). ρ represents Spearman correlation between module scores and fetal TEC expression indices, with * as p < 0.05. NES, normalized enrichment score.

We first evaluated for differences between naïve CD8^+^ T cells targeting TAAs with high versus low expression in fetal TECs (TAA^High-FT^ versus TAA^Low-FT^) in the unstimulated state. Naïve CD8^+^ T cells are largely quiescent with a homeostatic transcriptome, thus we utilized gene set enrichment analysis (GSEA) to assess coordinated pathway-level shifts. Naïve CD8⁺ T cells targeting TAA^High-FT^ showed increased expression in IFNα response signaling, mirroring findings in mice that naïve self-reactive CD8⁺ T cells display heightened type I interferon responsiveness.^32^ Additionally, naïve CD8⁺ T cells targeting TAA^High-FT^ showed increased expression of oxidative phosphorylation and MYC signaling pathways, but reduced IL-2-STAT5 and TNF-α/NF-κB signaling (Fig 4C and Extended Data Fig 4B). To test whether these pathways scaled with the TAA fetal TEC expression index, we used the leading-edge genes from each GSEA result to calculate corresponding module scores (Fig 4D). These pathway scores correlated with fetal TEC expression index (up to ρ = 0.77), consistent with a graded imprint of a hypofunctional programming from thymic exposure. Importantly, module scores from unsorted naïve CD8^+^ T cells, which are expected to primarily target foreign antigens (e.g. viral antigens not expressed in the thymus), most closely resembled those T cells specific for TAA^Low-FT^, consistent with a link between limited thymic exposure and heightened activation potential (Fig 4D). Together, these findings suggest that CD8⁺ T cells encountering high antigen levels in the fetal thymus are metabolically skewed toward quiescence (oxidative phosphorylation) and sensitivity to type I IFN, and away from signaling programs associated with robust effector activation (IL2-STAT5 and TNFα-NF-κB).

We next examined how naïve CD8^+^ T cells targeting TAA^High-FT^ versus TAA^Low-FT^ differentially responded to peptide stimulation. Recently activated naïve CD8⁺ T cells targeting TAA^Low-FT^ showed greater enrichment of activation-associated programs, including IFNγ responses and mTOR signaling (Fig 4E and Extended Data Fig 4C). In contrast, cells targeting TAA^High-FT^ showed comparatively muted enrichment of these pathways. This response followed a threshold pattern rather than a continuous gradient (Fig 4F), suggesting that only cells experiencing sufficient signaling strength can cross the threshold needed to overcome thymically imprinted tolerance.

At the gene level, naïve CD8⁺ T cells targeting TAA^Low-FT^ upregulated SRGN, consistent with acquisition of cytolytic effector functions (Fig 4G).^33^ In contrast, CD8^+^ T cells targeting TAA^High-FT^ retained elevated KLF2 expression, a transcription factor associated with naïve and quiescent states.^34^ Unexpectedly, CD8^+^ T cells targeting TAA^High-FT^ also showed reduced expression of LCP2 and FYB1, encoding the adaptor proteins SLP-76 and ADAP, which are critical mediators of proximal TCR signal transduction (Fig 4H). These TCR adaptor proteins are constitutively expressed in T cells and regulated post-translationally via rapid phosphorylation, rather than changes in mRNA abundance.^35,36^ Thus their transcriptional downregulation suggests a thymically imprinted attenuation of TCR signaling capacity. This reduced adaptor expression may contribute to the higher activation threshold observed with CD8^+^ T cells targeting TAA^High-FT^, and suggests a molecular mechanism by which fetal thymic exposure enforces durable hypofunctional programming in the surviving repertoire.

### Incorporating fetal thymic expression into TAA target selection enhances immunotherapeutic development

Having established a framework where fetal thymic antigen exposure shapes the functional potential of TAA-specific CD8⁺ T cells, we next asked whether this concept could help guide development of more effective immunotherapies. Many strategies under active investigation depend on the intrinsic immunogenicity of selected TAAs, including vaccines and adoptive transfer of peptide-expanded T cells. Even the identification of candidate receptors for engineered TCR-based therapies can be impacted by intrinsic TAA immunogenicity, which can constrain the ability to isolate functional TCRs. We reasoned that incorporating fetal TEC expression into target selection might enhance the likelihood of identifying antigens capable of eliciting effective responses.

Neuroblastoma is the most common extracranial solid tumor of childhood, and remains lethal in many patients despite intensive cytotoxic chemotherapy. Numerous efforts have focused on developing novel immunotherapies, though these tumors are characterized by low mutational burdens that leave few opportunities for neoantigen-directed strategies.^5^ Instead, greater emphasis has been placed on targeting non-mutated TAAs. A recent study highlighted PHOX2B, an oncofetal antigen with high tumor expression and minimal postnatal healthy tissue expression, as a promising candidate.^37^ Despite this favorable therapeutic window, repeated attempts to isolate high-affinity TCRs against PHOX2B have been unsuccessful. Notably, we find that while PHOX2B is not expressed by postnatal TECs, it is highly expressed by fetal TECs (Extended Data Fig 5A), suggesting that tolerance imparted by the fetal thymus may underlie its limited immunogenicity and hinder development of PHOX2B as an immunotherapeutic target.

To test whether incorporating fetal TEC expression could refine target prioritization in this setting, we reanalyzed the published neuroblastoma dataset.^37^ Of 4,210 candidate TAAs identified from cell lines or patient-derived xenografts, 161 had low healthy tissue expression, and only 11 demonstrated low fetal TEC expression (<0.5; Fig 5A). To evaluate their immunogenic potential, we selected five TAAs with the lowest fetal TEC expression (MAGE-B2, TSPY4, UPP2, MAGE-E2, and SIGLEC12) and compared them to five TAAs with high fetal TEC expression and with matched healthy tissue expression (PHOX2B, ZNF593, KRT9, SLC35E3, and DCX; Fig 5B and Supplemental Fig 5B). In peptide-stimulated PBMC assays, TAAs with low fetal TEC expression indices were more likely to elicit strong IFN-γ responses than those with high fetal TEC indices, demonstrating that fetal TEC expression can prospectively enrich for more immunogenic TAA targets (Fig 5C).

**Fig 5:**
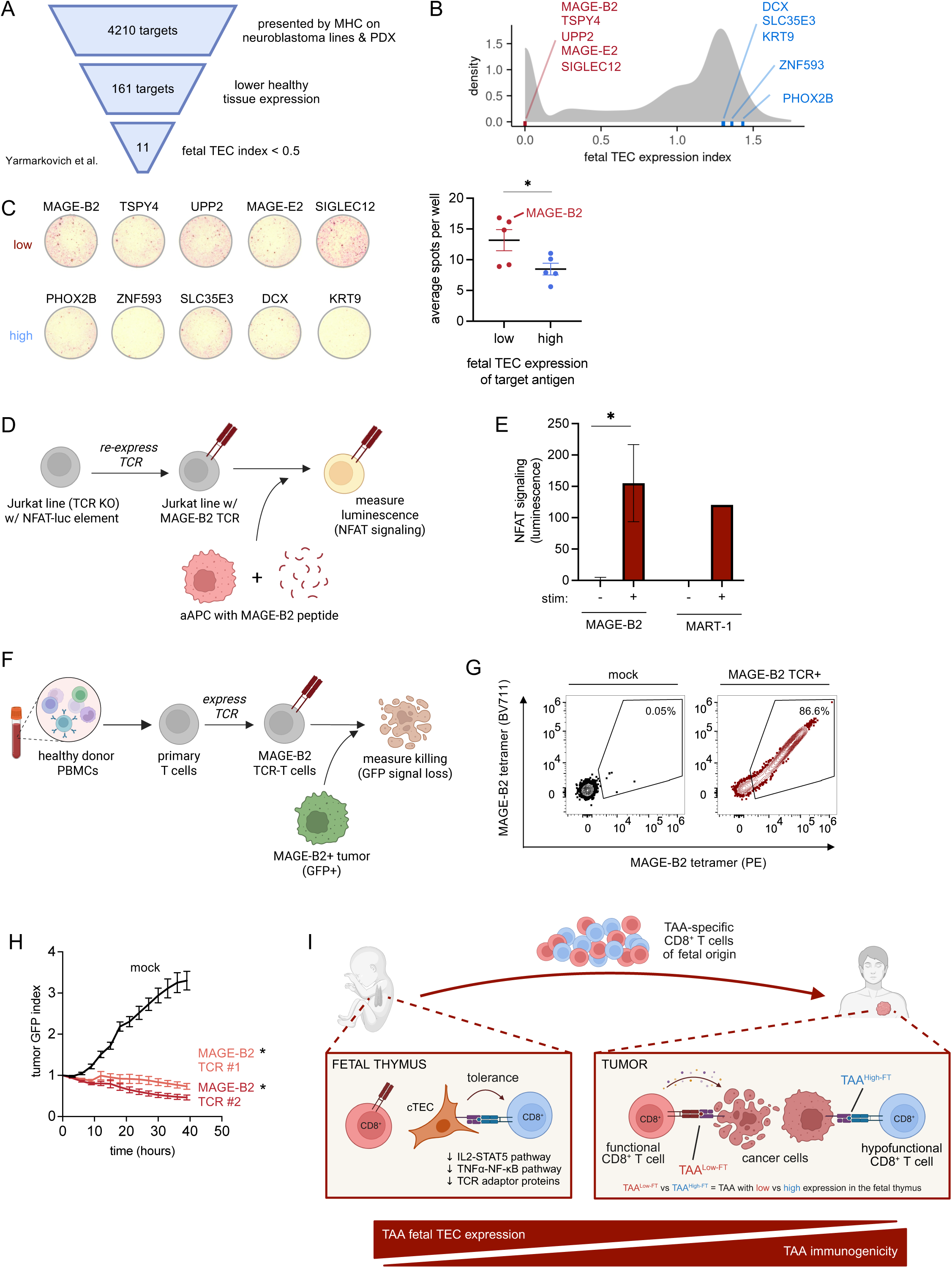
Incorporating fetal thymic expression into TAA target selection enhances immunotherapeutic development. **(A)** Reprioritization of TAA candidates from previously published immunopeptidomics data for neuroblastoma.^37^ Candidate targets were first identified based on documented presentation by MHC on neuroblastoma cell lines and patient derived xenograft (PDX) samples, then filtered for low healthy tissue expression (GTEx reference), and ranked by fetal TEC expression indices to exclude antigens with high thymic expression. This approach highlights how tolerance-informed filtering can refine therapeutic antigen selection. **(B)** Fetal TEC expression indices of selected candidate neuroblastoma TAAs. Five candidate TAAs with the lowest fetal TEC expression (red) and five with high fetal TEC expression (blue) are shown, overlying the distribution of fetal TEC expression indices for all transcribed genes in the genome, and illustrating broad variability in thymic expression among candidate targets. **(C)** Immunogenicity of neuroblastoma TAAs with low versus high fetal TEC expression indices, as assessed by IFNγ ELISPOT. Healthy donor HLA-A*02^+^ PBMCs were stimulated with pools of 5 peptides predicted to bind to HLA-A*02 (NetMHCpan) for 7 days, then repeat peptide pulsed for 24 hours prior to evaluation. Left, representative membranes; right, quantification of spots across n = 4 donors, showing that candidate targets with low fetal TEC expression indices were more likely to elicit a stronger IFNγ response than those with high fetal TEC expression. **(D)** Schematic of evaluating MAGE-B2 TCR signaling strength. MAGE-B2 TCRs were transduced into a Jurkat TCR^KO^ cell line with an NFAT-luciferase reporter element. Jurkat TCR^MAGE-B2^ cell lines were then challenged with peptide-pulsed aAPCs, and TCR-induced NFAT signaling measured by luciferase luminescence. **(E)** Strength of TCR-induced NFAT signaling for MAGE-B2 TCRs (n = 9) versus MART-1 TCR, a TAA noted to be highly immunogenic in Fig 1 (stimulated with MART-1 peptide). Results show MAGE-B2 TCRs induce strong NFAT signaling with peptide stimulation, comparable to that of MART-1 TCRs. **(F)** Schematic of evaluating the anti-tumor potential of MAGE-B2 TCRs. Healthy donor T cells were transduced to express MAGE-B2 TCRs and co-cultured with GFP⁺ MAGE-B2⁺ K562 tumor cells. Tumor killing was monitored over time by measuring GFP signal loss. **(G)** Transduction efficiency of MAGE-B2 TCR-T cells, measured using MAGE-B2 tetramers. Results show high levels of transduction in the MAGE-B2 TCR-T cells that are used in subsequent in vitro killing assays. **(H)** In vitro anti-tumor efficacy of MAGE-B2 TCR-T cells, compared to mock transduced T cells, showing potent killing of MAGE-B2^+^ HLA-A*02^+^ K562 tumors (endogenous MAGE-B2 expression). This strong antitumor response is consistent with the high immunogenic potential of TAA targets with low fetal TEC expression. **(I)** Schematic illustrating a model wherein a large fraction of TAA-specific T cells are fetal in origin, educated by fetal TECs, and thereby imparted with programming that shapes their immunogenic potential in the periphery. Thus, incorporating fetal thymic expression patterns into antigen selection can rationally prioritize TAAs with greater inherent immunogenic potential for therapeutic development. * represents p < 0.05 by t test (for h, performed at hour 40)

Among the neuroblastoma TAAs with low fetal TEC expression, we selected MAGE-B2 for further study. In contrast to PHOX2B, highly-effective TCRs against MAGE-B2 can be readily isolated from peptide-stimulated PBMCs.^38–40^ MAGE-B2 TCRs triggered robust NFAT activation when introduced into Jurkat reporter cells and stimulated with peptide, at levels comparable to those seen with MART-1 TCRs, a well-established immunogenic control from our initial studies (Fig 5D-E). Importantly, this increased signaling strength translated into meaningful antitumor activity. When transduced into primary T cells, MAGE-B2-specific TCRs mediated potent killing of MAGE-B2^+^ tumors in co-culture assays (Fig 5F-H). Thus, MAGE-B2 exemplifies the value of incorporating fetal TEC expression into TAA selection, providing a proof-of-concept that this framework prospectively helps identify antigens with higher intrinsic immunogenicity and greater translational potential.

## DISCUSSION

TAAs have long represented an attractive class of targets for cancer immunotherapies, offering the potential to expand beyond the limited and highly variable pool of mutationally-derived neoantigens. The impressive responses achieved by targeting TAAs such as MAGE-A4 and NY-ESO-1 demonstrate that self-antigen immunity can be therapeutically harnessed.^8–11^ Yet, most TAAs fail to elicit robust antitumor immunity, and the basis for this variability has remained poorly understood.

Here, we show that despite TAAs all being self-antigens, they display a striking variability in immunogenicity that appears to be shaped by thymic tolerance established early in human development. We find that TAA expression by fetal TECs is a strong predictor of its immunogenicity in both *in vitro* T cell stimulation assays and *in vivo* patient responses to checkpoint blockade. TAAs that are highly expressed by fetal TECs are associated with CD8^+^ T cells with a hypofunctional programming and reduced effector potential. Together, these data support a model in which a large fraction of TAA-specific T cells are fetal in origin, educated by fetal TECs, and thereby imparted with programming that shapes their immunogenic potential in the periphery (Fig 5I).

That TAAs could have variable immunogenicity based on fetal thymic expression was unexpected but is consistent with prior observations in human immunology. A substantial fraction of the human T cell repertoire is generated prenatally. Many infants who undergo thymectomy for congenital heart disease on the first days of life maintain relatively normal T cell numbers and diversity for decades, underscoring the persistence and breadth of fetally derived clones.^41^ Consistent with this, we find that TAA-specific TCRs share structural features characteristic of fetal repertoires, including shorter CDR3 regions, further supporting a prenatal origin. Fetally derived T cells have also been implicated in autoimmunity, with cord blood being enriched for T cells recognizing self-antigens,^25^ and mouse studies have suggested that exposure to the fetal thymus is required for the development of autoimmune diabetes.^42^ Our findings extend these observations, linking fetal thymic imprinting to the regulation of TAA reactivity.

Mechanistically, our data indicate that fetal TEC-mediated tolerance of CD8^+^ T cells to TAAs extends beyond clonal deletion, with evidence pointing to a predominant role for cTECs rather than the more well studied mTECs. This finding aligns with prior work demonstrating that self-reactive T cells often escape thymic deletion yet remain functionally restrained.^14,30,31,43^ Across both human and murine systems, self-specific T cells persist at near-normal frequencies, but exhibit elevated activation thresholds and require strong inflammatory cues for stimulation. By uniquely focusing on naïve TAA-specific CD8⁺ T cells, our study shows that this state is thymically imprinted rather than peripherally acquired. Exposure to TAAs in the fetal thymus appears to confer a distinct transcriptional, signaling, and metabolic state on developing CD8⁺ T cells, characterized by altered TCR scaffold components, increased sensitivity to type I interferons,^32^ and a metabolic profile consistent with lower effector potential. Together, these findings suggest that prenatal thymic antigen exposure imprints a spectrum of functional restraint within the human CD8⁺ T cell repertoire, paralleling thymic tuning mechanisms previously described for CD4⁺ T cells.^44^ Validation of this model will require additional experimental studies, which are the focus of our ongoing work. Further, while our findings highlight a thymic, fetal component to TAA-specific tolerance, peripheral mechanisms of tolerance such as anergy and exhaustion are also likely to modulate these responses. These processes may act in concert with thymic fetal imprinting to further shape the magnitude and durability of antitumor immunity.

This framework also provides new context for understanding why some tumors respond to immune checkpoint inhibitors while others do not. Although response rates correlate with mutational burden and neoantigen load, these features alone fail to explain the high variability seen across cancers and patients. Recent studies have suggested that immunity to non-mutated cancer antigens contributes to responses to checkpoint inhibitors, and our findings provide evidence supporting this idea.^7^ We show here that TAAs that “drop out” under immune checkpoint therapy tend to be poorly expressed in the fetal thymus, suggesting that reduced fetal exposure allows CD8⁺ T cells targeting these antigens to escape thymic tolerance and mount effective antitumor responses.

Furthermore, these findings suggest that fetal thymic expression serves as a previously unrecognized filter helpful in determining which TAAs are viable immunotherapy targets. Indeed, the most successful TAA-targeted therapies, such as MAGE-A4 and NY-ESO-1, are directed against antigens with minimal fetal thymic expression.^8–11^ In contrast, targeting TAAs like WT1, PRAME, and Survivin, which are highly expressed during fetal thymic development, have shown only modest efficacy or require minimal residual disease settings to demonstrate any benefit.^19,45,46^ Incorporating fetal thymic expression patterns into antigen selection could therefore rationally prioritize TAAs with greater inherent immunogenic potential.

To illustrate this principle, we reanalyzed published neuroblastoma immunopeptidomic data in which the original authors prioritized PHOX2B, a canonical neuroblastoma antigen, for TCR discovery through peptide stimulation of PBMCs.^37^ These efforts failed to expand antigen-specific TCRs for sequencing, a result consistent with our finding that PHOX2B is highly expressed by fetal TECs. Interestingly, PHOX2B is not expressed in the postnatal thymus, underscoring the importance of considering the fetal contribution of thymic tolerance, in particular, when evaluating TAA immunogenicity. In contrast, our framework reprioritized MAGE-B2, which shows low fetal thymic expression and correspondingly higher immunogenicity in preclinical models. These findings highlight how incorporating fetal thymic expression data into antigen discovery can refine target selection and enhance immunotherapeutic development, which may include vaccine design, adoptive transfer of peptide-expanded T cells, and identification of antigen-specific TCRs for engineering of TCR-transduced T cells.

In summary, our findings reveal that thymic tolerance, particularly that established during fetal development, constrains the intrinsic immunogenic potential of TAAs by imprinting a hypofunctional state on CD8⁺ T cells recognizing those antigens. This developmental aspect of thymic tolerance helps explain the variability in TAA immunogenicity and provides a framework for selecting antigens that would have the best chance of being clinically effective. More broadly, these results suggest that for cancers with low neoantigen burdens, including hematologic and pediatric malignancies, effective next-generation therapies may require not only prioritizing TAAs over neoantigens but also harnessing distinct T cell populations with unique functional programming shaped by fetal rather than postnatal thymic selection.

## METHODS

### Cells and culture conditions

Human PBMCs from healthy donors were obtained from Leukocyte Reduction System (LRS) chambers collected during platelet donation and purchased from the Stanford Blood Center under an IRB-exempt protocol. All donors were adult (ages 37-89 yo), HLA-A*02 positive and CMV seronegative. PBMCs or CD8⁺ T cells were isolated using Lymphoprep density gradients (StemCell Technologies) or CD8⁺ T Cell Isolation Kits (Miltenyi), respectively, and cryopreserved in CryoStor CS10 (Lonza) for later use.

With the exception of primary T cells transduced with exogenous TCRs, PBMCs and T cells were cultured in RPMI 1640 (Gibco) supplemented with 10% FBS (R&D Systems), 100 U/mL Penicillin/Streptomycin (Gibco), 100 µg/mL Normocin (Invivogen), 1x MEM non-essential amino acids (Gibco), 1 mM sodium pyruvate (Gibco), 1x insulin-transferrin-selenium (Gibco), and 10 mM HEPES (Gibco). Primary T cells transduced with exogenous TCRs were cultured in AIM-V (Gibco) supplemented with 5% FBS, 100 U/mL Penicillin/Streptomycin, and 10 mM HEPES. Media was exchanged every 2-3 days.

We previously reported generation of a Jurkat cell line with TCR KO expressing an NFAT-luciferase (J76-NFATRE-luc) and a K562-based artificial APC expressing HLA-A*02.^55^ These cells were maintained in RPMI 1640 supplemented with 10% FBS, 100 U/mL Penicillin/Streptomycin, and 10 mM HEPES. LentiX 293T and 293GP cell lines were maintained in DMEM (Gibco) supplemented with 10% FBS, 100 U/mL Penicillin/Streptomycin, and 10 mM HEPES.

### PBMC peptide stimulation

PBMCs were cultured at a density of 6 x 10^6^ / mL and stimulated with 1 μg/mL of peptide (Elim Bio and GenScript) dissolved in DMSO. Matching volumes of DMSO were used as controls. Media was exchanged every 2-3 days and additional peptide was not reintroduced unless otherwise noted. PBMCs remained in culture for 7 days, then harvested for subsequent analyses.

### T cell functional assays

CellTrace Violet proliferation assays were conducted by staining PBMCs immediately prior to peptide stimulation. PBMCs were stained with 5 μM CellTrace Violet (Invitrogen) at a concentration of 10^7^ cells / mL per manufacturer’s recommendations. Proliferation was then evaluated by flow cytometry after 7 days of culture. Upregulation of activation markers CD25 and CD38 was evaluated 7 days after initial stimulation, following peptide rechallenge 24 hours prior to assessment. CD107a degranulation assays were conducted by flow cytometry 7 days after initial peptide stimulation. Six hours prior to assessment, cultures were rechallenged with peptide in the presence of 2 μM monensin and a CD107a antibody. All T cell functional assays were evaluated on antigen-specific (tetramer-positive), activated (CD38^+^) cells.

### Generation of peptide-MHC tetramers

Peptide-MHC tetramers were generated as previously described.^56^ In brief, the HLA-A*02 α-chain and β2m proteins were overexpressed separately in BL21 DE3 *E. coli* (Invitrogen) and isolated from inclusion bodies. Monomers were refolded in the presence of the UV cleavable peptide (KILGFVFJV; J = 3-amino-3-(2-nitrophenyl)propionic acid residue; Elim Bio), biotinylated, concentrated, and stored with 20% glycerol at −80°C. Exchange reactions were subsequently performed with 100 μg/mL monomer and 200 μM of peptide in PBS (pH 7.4) by exposing the mixture to 365 nm UV light for 30 minutes on ice using a UV Stratalinker 2400 (Stratagene). Plates were then incubated at 37°C for 30 minutes to complete the exchange.

Tetramers were subsequently formed by incubating the exchanged monomer with streptavidins (Biolegend) tagged with fluorophores with or without oligo-barcodes (TotalSeq C; Biolegend) for 30 minutes (Supplemental Table 1). Excess streptavidin was quenched by the addition of 0.4 mM biotin. Tetramers were stored up to 2 weeks at 4°C, and spun at 13,000 g for 10 min at 4°C to remove aggregates immediately prior to use.

### Flow cytometry and cell sorting

All samples were analyzed with a Penteon (Agilent) or flow sorted with a FACSAria Fusion (BD). Tetramer staining was performed for 60 minutes on ice unless otherwise noted. TAA tetramers were stained in pairs, generated with PE– and BV711-streptavidins. Samples were simultaneously stained with CMV pp65 tetramers generated with APC– and PE-Cy7-streptavidins as a negative control. TAA-specific T cells were defined as double negative for CMV tetramers and double positive for TAA tetramers to ensure specificity. Following tetramer staining, the following were used for T cell phenotyping: Fixable Live/Dead Blue (or Fixable Live/Dead Near IR for CellTrace Violet proliferation experiments, Invitrogen), CD3 BV605 (Biolegend clone SK7), CD4 FITC (Biolegend clone RPA-T4), CD8 BV785 (Biolegend clone RPA-T8), CD38 BV510 (Biolegend clone HIT2), CD107a APC-Cy7 (Biolegend clone H4A3) and CD25 BUV737 (BD clone 2A3). All samples were stained in the presence of Human Fc block (Biolegend) and Brilliant Stain Buffer (BD). Example gating shown in Supplemental Fig 1.

### TEC atlas scRNAseq dataset analysis

#### Quality control, integration, and cell annotation

Previously published single-cell RNAseq data from the human thymic atlas were downloaded from Zenodo (https://zenodo.org/records/3711134) as a raw count matrix (h5ad format) with associated metadata.^22^ Analyses were performed in R (v4.2.2) using Seurat (v4.4.0).^57^ Thymic epithelial cell (TEC) samples were subsetted from thymocyte samples based on metadata and filtered to retain cells expressing 500 – 6,000 genes, <40,000 UMIs, and <20% mitochondrial transcripts.

Donor-specific genes (TRAV, TRBV, IGH, IGK/L) and mitochondrial genes were removed from the counts matrix, whereas HLA genes were retained given their importance in distinguishing cortical (cTEC) and medullary (mTEC) subtypes. Data were normalized using SCTransform with regression of cell-cycle effects, and the top 3,000 highly variable genes were selected for downstream analyses. Batch effects across samples were corrected using Harmony^58^ (v1.2.3; kmeans.nstart = 100, kmeans.iter.max = 5,000). Dimensionality was assessed by JackStraw plots, and the top 20 Harmony components were used for UMAP projection (n.neighbors = 20, min.dist = 0.5).

Unbiased clustering was performed with FindNeighbors and FindClusters (resolution = 0.2). Clusters were annotated as cTEC or mTEC subtypes based on canonical markers: cTECs (PSMB11^hi^), mTECs (PSMB11^lo^ PSMB9^hi^), and further mTEC subsets defined as: (I) KRT14^hi^, (II) AIRE⁺/FEZF2⁺, (III) KRT10^hi^, (myoid) MYOG⁺/MYOD1⁺, and (neuroendocrine) NEUROD1⁺/NEUROG1⁺.

#### Calculating TEC expression of TAAs (pseudobulk and TEC expression index)

Following quality control and appropriate cell annotation, HLA genes were removed from the matrices and per cell data normalized using Seurat’s NormalizeData function. TAA pseudobulk expression across all TECs was calculated per gene as the mean of all expression levels. To generate TEC expression indices, the expression matrix was ranked on a per-gene basis, and the mean expression of the top expressors (0.01% to 100%) was calculated. Aggregated TEC expression values were transformed as log_10_(TEC expression + 0.1) + 1 to accommodate zero values and stabilize variance at the aggregate level. This process was repeated as needed on subsetted versions of the normalized data matrix based on cell annotations (i.e. developmental state or cTEC vs. mTEC). To control for differences in sequencing depth and cellular representation across samples, these analyses were repeated on a downsampled dataset. UMI counts for infant and fetal groups were probabilistically downsampled to the median UMI count of adult samples (developmental group with lowest average UMI count) per cell using binomial thinning, preserving the relative distribution of gene expression while equalizing total counts. Cell numbers were then downsampled for adult and fetal groups to match the infant group (the developmental group with the fewest cells).

Linear regression (lm function) was used to evaluate associations between TEC expression and the frequencies of tetramer-positive CD8⁺ T cells after stimulation. Analyses were performed using either pseudobulk TEC expression values or TEC expression indices. For regression modeling, frequencies of tetramer-positive CD8⁺ T cells were standardized within donors (z-score), shifted to positive values to enable log transformation, and log_10_ transformed to compress dynamic range and improve linearity.

### TAA dropout in melanoma datasets

#### FASTQ data accessing

RNAseq data from three previously published datasets for melanoma patients receiving checkpoint inhibitor treatment were analyzed.^27–29^ Each publication had paired pre-treatment and on-treatment samples for analysis. FASTQ files were downloaded from the Sequence Read Archive (SRA) using sratoolkit (v3.2.0; Supplemental Table 2). Individual runs were retrieved with prefetch, and FASTQ files were generated using fasterq-dump with the ––split-files option to separate read pairs.

#### Inferring class I HLA typing

Each subject’s class I HLA typing was inferred using OptiType (v1.0).^53^ Reads aligning to HLA loci were first extracted from FASTQ files using *razers3* (v3.5.8)^52^ with parameters –i 95 –m 1 –dr 0 against a custom HLA reference (Optitype Github). The resulting alignments were converted back to FASTQ format using samtools (v1.3.1)^51^ for downstream HLA typing with OptiType executed with default parameters.

#### TPM counts table generation

Processed RNAseq count tables for Riaz et al.^27^ and Du et al.^28^ were downloaded from the Gene Expression Omnibus (GEO; accession numbers GSE91061 and GSE168204). Gide et al.^29^ data were processed to generate a TPM counts table. Raw paired-end sequencing reads were trimmed using fastp (v0.12.4)^50^ to remove adapter sequences and low-quality bases. Read quality after trimming was assessed using FastQC (v0.11.8). Trimmed reads were aligned to the human reference genome (GRCh38) using STAR (v2.7.10b)^48^ with default parameters, outputting coordinate-sorted BAM files. The STAR genome index was generated from the GRCh38 reference assembly and GENCODE v36 gene annotation. Alignments were filtered to retain high-quality reads (MAPQ ≥10), excluding unmapped, secondary, and supplementary alignments, using samtools (v1.3.1), and indexed for downstream analysis. Gene-level counts were then quantified using HTSeq (v0.11.2)^49^ with non-stranded read counting. Raw gene counts were then converted to transcripts per million (TPM) by normalizing for both sequencing depth and gene length, where gene lengths were defined as the sum of non-overlapping exon lengths per gene from the same GENCODE v36 annotation.

#### Predicting presented TAA epitopes

For each dataset, subject, and timepoint, TAA candidates were identified by comparing tumor gene expression (TPM) to median expression levels across healthy tissues from the Genotype-Tissue Expression (GTEx) Project, version 10.^59^ Genes were classified as putative TAAs if their expression was <5 TPM in all healthy tissues and at least 50-fold higher in tumor samples relative to the corresponding healthy median. Expression in testis was excluded from this analysis given its immunologically privileged status.

Protein sequences for each TAA were retrieved using biomaRt (version 2.58.2; useEnsembl, Ensembl release 105), and canonical isoforms were identified by cross-referencing annotated gene IDs with AnnotationHub (version 3.10.1). For each subject, epitopes derived from expressed TAAs were predicted based on individual HLA genotypes using NetMHCpan (v4.1b).^54^

#### Evaluating TAA dropout features with checkpoint blockade response

To assess features associated with TAA dropout during checkpoint inhibitor therapy, predicted TAA-derived epitopes were quantified in pre– and on-treatment tumor samples. Epitopes predicted to bind multiple HLA class I alleles were weighted according to the number of presenting alleles. TAAs detected pre-treatment but absent on-treatment were classified as dropped, whereas those still detected were classified as retained. Each epitope was assigned a TEC expression index based on its parent TAA. For each patient, mean TEC expression indices were calculated separately for dropped and retained epitopes. ROC analyses were performed across all patients to evaluate which TEC expression index most effectively distinguished mean of dropped or retained epitopes per subject. AUC values were compared across TEC developmental subsets.

### TAA-specific CD8^+^ T cell multimodal profiling

#### Library preparation, sequencing, and alignment

Paired unstimulated and stimulated CD8^+^ T cells from the same donors (n = 12) were used. Stimulated cells were generated by PBMC peptide stimulation, as above. Seven days after stimulation, CD8^+^ T cells were isolated from the cultures using a CD8^+^ T cell Isolation Kit (Miltenyi). Unstimulated CD8^+^ T cells were isolated from PBMCs prior to cryopreservation and were thawed the day of analysis. Isolated CD8^+^ cells were stained with oligo-tagged tetramers for 60 minutes at 37°C in the presence of Fc block and Monocyte Block (Biolegend). To minimize the possibility of tetramer binding inducing TCR signaling, all staining and processing was performed in the presence of 50 μM dasatinib (Stem Cell Technologies).^60^ Two tetramers with different oligo-barcodes were used per TAA to increase specificity. Staining with fluorescently tagged antibodies for CD3, CD4 and CD8, as well as oligo-tagged hashtag antibodies for donor identification, was then performed. All donor samples were then combined and stained with oligo-tagged antibodies for cell surface protein analysis (Biolegend TotalSeq-C Human Universal Cocktail, V1.0; stained at ∼7.5 x 10^7^ per test). Tetramer positive cells were then flow sorted and cell capture performed on a 10x Genomics Chromium Controller targeting 10,000 cells per lane, with 6-7 lanes per stimulation condition. Unsorted cells for both conditions were also run in a separate lane. Individual gene expression (GEX), Antibody Derived Tag (ADT), and T Cell Receptor (TCR) libraries were prepared according to the Chromium Next GEM Single Cell 5’ Reagents Kit v2 protocol (10x Genomics). Final libraries were sequenced on an Illumina Novaseq6000 using 151/10/10/151 base pair configuration (Novogene). Demultiplexed FASTQ files were aligned to the hg38 reference genome (GRCh38-2020-A) and processed using the CellRanger (7.0.0) multi command.

#### Quality control, data integration, and cell annotation

All analyses were performed in R (v4.2.2) using Seurat (v4.4.0). Cells expressing >200 genes, >2,000 UMIs, and <10% mitochondrial transcripts were retained. Surface protein measurements (hashtags, tetramers, and antibody-derived tags) were normalized using centered log-ratio transformation. Donor identities were assigned with HTODemux (positive.quantile = 0.9999), and doublets excluded (identified as cells with >1 donor call).

Tetramer specificity thresholds were defined by manual inspection (Extended Data Fig 1B) at the boundary of the negative population. Cells were designated TAA-specific if double-positive for both cognate tetramers and negative for all others. To minimize spurious assignments, only TAAs with tetramer concordance > 0.5 and detectable TAA-specific cells in ≥ 25% of donors were retained for analysis (Extended Data Fig 1C-D). Tetramer sorted cells without a positive tetramer call were excluded. Donor-specific genes (TRAV, TRBV, IGH, IGK/L, HLA) and mitochondrial genes were removed for downstream analyses.

Unstimulated and stimulated datasets were normalized independently using SCTransform with regression of cell-cycle effects, merged, and then integrated with Harmony by sample and stimulation condition (v1.2.3; kmeans.nstart = 100, iter.max = 5,000). Dimensionality was assessed by JackStraw plots, and the top 40 Harmony components used for UMAP projection (n.neighbors = 20, min.dist = 0.2). Unbiased clustering (FindNeighbors, FindClusters; resolution = 1.4) identified contaminating CD19⁺ B cells, CD4⁺ T cells, and myeloid cells, which were excluded. SCTransformation, Harmony integration, UMAP, and clustering were then repeated (dims = 1:15, resolution = 0.4).

Memory states were annotated from canonical markers at the protein and RNA levels: naïve (CD45RA⁺ CD45RO⁻ CD62L⁺), central memory (CD45RA⁻ CD45RO⁺ CD62L⁺), effector memory (CD45RA⁻ CD45RO⁺ CD62L⁻), activated memory (CD45RA⁻ CD45RO⁺ CD69⁺), and exhausted (CD45RA⁻ CD45RO⁺ PD-1⁺ 2B4⁺ TIGIT⁺). Activation intensity within the naïve subset was further stratified as low, mid, or high (CD69, KI67, GZMA).

#### Transcriptional and pathway analysis

Naïve CD8⁺ T cells were isolated from the fully annotated dataset and analyzed separately by stimulation condition. For each condition, data were re-normalized using SCTransform, re-integrated with Harmony, and visualized by UMAP (dims = 1:15, n.neighbors = 10, min.dist = 0.005). Contaminating memory (CD45RA⁻ CD45RO⁺) cells were excluded, and for the unstimulated dataset, recently activated cells (CD69⁺) were removed to minimize effects of peripheral antigen exposure. These filtered, condition-specific datasets were then re-normalized using SCTransform and used for differential gene expression and GSEA analyses.

To assess transcriptional differences between CD8^+^ T cells targeting TAAs that are differentially expressed in the fetal thymus, we stratified cells based on their antigen’s fetal TEC expression index (low <0.75; high >0.75) and calculated differential gene expression using Seurat’s FindMarkers (Wilcoxon test, SCT data slot). Ranked gene lists were analyzed by GSEA using fgseaMultilevel (fgsea v1.24.0)^47^ with the MSigDB Hallmark v2023.1 gene sets (FDR < 0.1), followed by collapsing of related pathways (collapsePathways).

To validate and quantify the continuous relationship between pathway activity and target TAA fetal thymus expression, module scores were computed from leading-edge genes for each significant GSEA pathway (AddModuleScore). Spearman correlations were then calculated between pathway module scores and the fetal TEC expression index. For visualization, cells were binned into low (<0.5), mid (0.5 – 1.1), and high (>1.1) groups to illustrate graded trends. To identify specific genes associated with tolerance-related extremes, differential expression was performed on CD8^+^ T cells targeting TAAs with the lowest (<0.5) and highest (>1.1) fetal TEC expression indices, using Seurat’s FindMarkers (Wilcoxon test, SCT data slot).

#### Calculating TCRβ CDR3 lengths

Single-cell TCR sequencing data (CellRanger output “filtered_contig_annotations.csv”) were annotated with tetramer specificity and memory state using matching cell barcodes from the Seurat object. Clonotypes were identified using Immunarch (v0.9.0; repLoad function), and TCRβ CDR3 lengths were measured directly from the CDR3 nucleotide sequences.

### Reanalysis of neuroblastoma immunopeptidomics data

Candidate antigens previously reported to be presented by HLA on neuroblastoma cells lines or PDX models by mass spectrometry were collated from supplemental data.^37^ Targets were narrowed to those that had low healthy tissue expression (GTEx Project, version 10; TPM < 10),^59^ and then ranked by fetal TEC expression index. Five TAAs with the lowest fetal TEC expression index were chosen for subsequent analysis, as well as five TAAs with high fetal TEC expression indices and with matching peripheral tissue expression.

## ELISPOT

For each TAA evaluated, peptide pools (Elim Bio) were generated from the top 5 predicted binders to HLA-A*02 using NetMHCpan (v4.1b). PBMCs were pulsed with peptides (1 µg/mL per peptide) and cultured for 12 days, adding 20 U/mL interleukin-2 (IL2, Peprotec) on days 3, 5, and 7. PBMCs were then analyzed by IFNγ ELISPOT. 5 x 10^5^ cells were plated per well in 96-well ELISPOT plates coated with anti-IFNγ antibody (2 µg/mL; MabTech). PBMCs were then restimulated with peptides (1 µg/mL per peptide) and incubated for 24 hours. Spots were then revealed with anti-IFNγ biotinylated detection antibody (0.3 µg/mL; MabTech), ExtrAvidin-Alkaline Phosphatase (1:1,000 dilution; Sigma Aldrich), and BCIP/NBT (5-bromo-4-chloro-3-indolyl-phosphate/nitro-blue tetrazolium chloride; Sigma Aldrich). Spots were imaged and counted (positive if ≥ 75 μm) using an a Cytation 7 (Agilent).

### NFAT reporter assay for TCR signaling

#### Lentiviral vector cloning, supernatant production and transduction

Downstream signaling strength of MAGE-B2 TCRs using an NFAT-luciferase reporter gene was evaluated as previously described.^55^ In brief, CDR3 regions for each TCR (Supplemental Table 3) were synthesized as oligonucleotides (IDT), annealed, and inserted into the corresponding second-generation N103 lentiviral backbone from an in-house library containing all human V-genes and matched constant regions for each chain. Plasmids were amplified and purified from Stbl3 E. coli. Lentiviral supernatants were produced via transient transfection of the Lenti-X 293T line (Takara), as previously described.^55^ Lenti-X 293T cells were transfected via ViaFect (Promega) with the expression plasmids (both TCRα and TCRβ), as well as pMD2.G envelope and psPAX2 packaging plasmids. Viral supernatants were collected 48 h later, clarified, and concentrated with Lenti-X Concentrator (Takara). J76-NFATRE-luciferase Jurkat cells were transduced in 96-well plates by spinoculation with 8 μg/mL polybrene (Sigma Aldrich). TCR^+^ Jurkat cells were flow sorted for purity (CD3 and tetramer staining).

#### Jurkat-aAPC-peptide co-culture

TCR-transduced J76-NFATRE-luciferase Jurkat cells were co-cultured with HLA-A*02^+^ K562-based, artificial antigen presenting cells (aAPCs) in the presence of TAA peptides at 1 µg/mL. Cells were incubated at 37°C for 18 hours. NFAT activity was then read out as luminescence (BioTek Cytation 7 plate reader, Agilent), after adding Nano-Glo Luciferase Assay reagent (Promega #N1630) according to the manufacturer’s instructions.

### In vitro tumor killing assay

#### Retroviral vector cloning, supernatant production, and transduction

MAGE-B2 TCRs were cloned into the MSGV retroviral expression vector using homology-directed assembly (Twist Bioscience). TCRβ and TCRα chains were linked by a furin-T2A self-cleaving peptide sequence. Human TCR constant regions were replaced with murine constant regions, as previously described, to promote correct pairing of exogenous α and β chains and minimize mis-pairing with endogenous TCRs within primary human T cells.^61^

Retroviral supernatants were produced via transient transfection of the 293GP cell line, as described.^62^ Briefly, 293GP cells were transfected via Lipofectamine 2000 (Invitrogen) with the expression plasmids and plasmid for the RD114 envelope protein. Supernatants were collected 48 h after transfection. Monocyte depleted PBMCs were activated with anti-CD3/CD28 beads (Invitrogen) in a 3:1 bead:cell ratio with 40 IU/ml IL-2 (Peprotec) for 3 days. Activated T cells were then retrovirally transduced on days 3 and 4 using Retronectin (Takara) coated plates, and cultured in 300 IU/ml IL-2. Anti-CD3/ CD28 beads were removed on day 5. Media and IL-2 were changed every 2 days. Transduction efficiencies were measured by tetramer staining and were 60 – 90% for all MAGE-B2 TCRs. Killing assays were subsequently performed on day 9 after initial activation.

#### Incucyte killing assay

5 x 10^4^ GFP-positive tumor cells and 4 x 10^5^ MAGE-B2 TCR T cells (E:T ratio 8:1) were cocultured in 200 μL complete AIMV at 37°C for up to 72 hours. Plates were analyzed every 3 hours using the IncuCyte Live-Cell Analysis System (Sartorius). Four images per well at 10× zoom were collected at each time point. Total integrated GFP intensity per well was measured. Values were normalized to the starting measurement and plotted over time.

### Quantification and statistical analysis

FlowJo v.10.10.0 was used to analyze flow cytometry data. All statistical analyses were performed using R version 4.2.2 or GraphPad Prism 10. Individual statistical tests for various analyses are listed in their respective figure legends. All statistical tests were two-sided unless otherwise stated. Statistical significance is indicated by * (p < 0.05, or adjusted p < 0.05 when indicated), unless otherwise stated. There were no formal statistical tests to predetermine sample sizes, and investigators were not blinded to sample conditions during the analysis.

## DATA AVAILABILITY

The multimodal single-cell data generated from this study can be obtained from the NCBI GEO database with accession number GSE310276. The detailed information of all public datasets has been provided in Supplemental Table 1. Any additional information required to reanalyze the data reported in this paper is available from the lead contact upon request. Further information and requests for resources should be directed to the lead contact, Mark Davis.

## CODE AVAILABILITY

No new code packages were developed in this study.

## ACKNOWLEDGMENTS

We kindly thank Crystal Mackall (Stanford University) for the 293GP cell line, MSGV vector plasmid, and RD114 plasmid. Cell sorting was performed in the Stanford Shared FACS Facility on the FACSAria Fusion purchased by the Parker Institute for Cancer Immunotherapy. This work was supported by the National Institute of Health (NIH, U19 AI057229 and P01 AI153559). AHL’s contribution to this work was supported by the Stanford Maternal & Child Health Research Institute, Doris Duke Charitable Foundation, and the Hyundai Hope On Wheels. We thank all members of the Davis lab for critical input and discussion, and we thank Kara Davis (Stanford University) for her thoughtful review of this manuscript.

## AUTHOR INFORMATION

Conceptualization, AHL and MMD; Methodology, AHL; Software, AHL, VS, and NAB; Investigation, AHL, KRT, NMP, CB, TB; Data Curation, AHL and NAB; Writing – Original Draft, AHL; Writing – Review & Editing, AHL, KRT, NMP, CB, TB, NAB, MMD; Visualization, AHL; Supervision, AHL and MMD; Funding Acquisition, AHL and MMD.

## ETHICS DECLARATION

The authors declare there are no conflicts of interest.

**Extended Data Fig 1:**
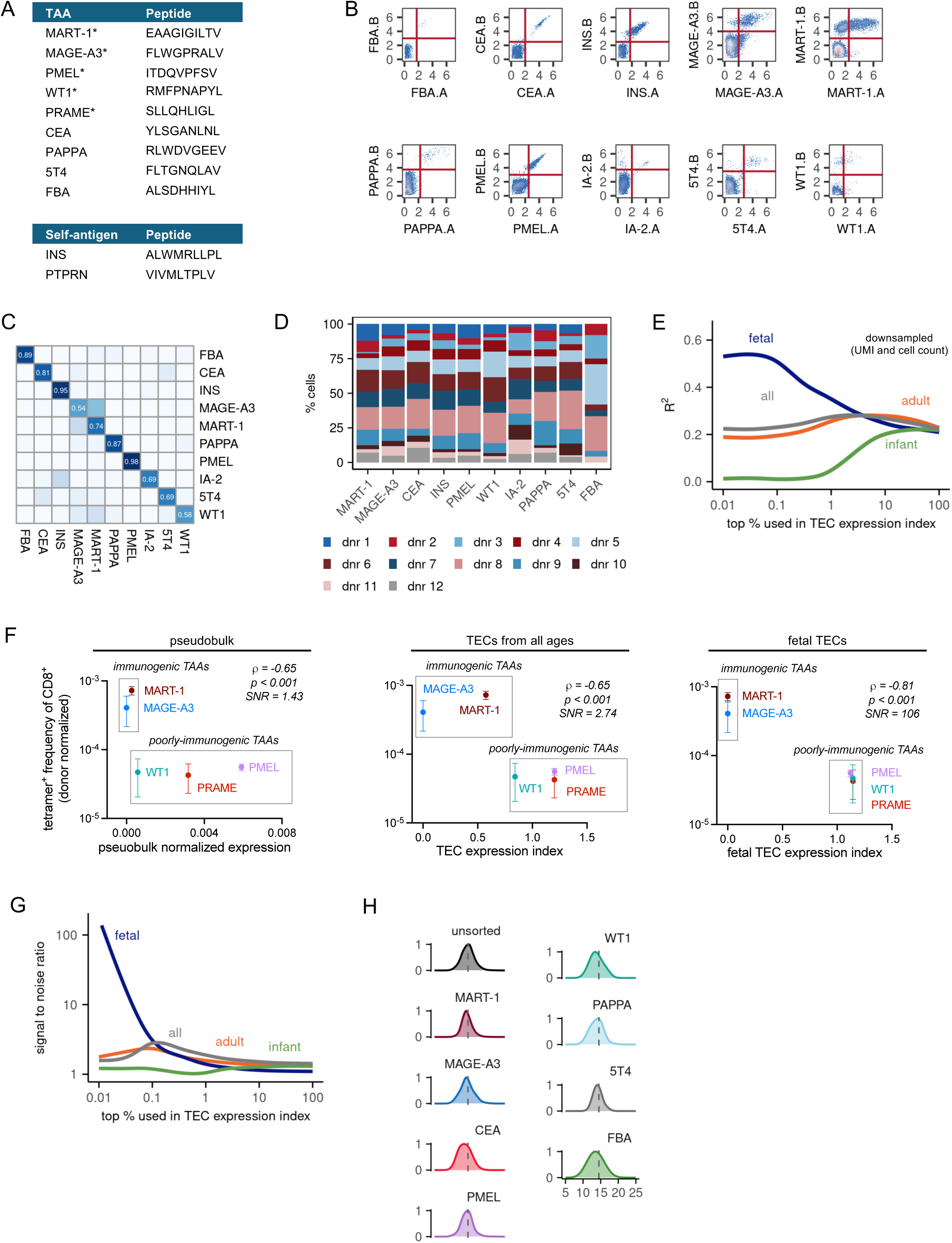
Quality control and validation of antigen specificity, TEC expression index robustness, and TCR features. **(A)** TAA and self-antigen peptide sequences used throughout *in vitro* stimulation assays. * denotes those used in Fig 1. All antigens were used in Fig 2. **(B)** Thresholds for assigning tetramer specificities using oligo-tagged tetramers and CITE seq. Each TAA-specificity had two unique oligo-tagged tetramers to enhance specificity. Plots represent normalized counts. Thresholds were set manually at boundary of negative population. **(C)** Concordance plot of tetramer specificities. Values indicate the frequency that a CD8^+^ T cell positive for one tetramer is also positive for the second. All specificities analyzed have high concordance, lending confidence to the tetramer calls. **(D)** Donor representation among all tetramer positive CD8^+^ T cells, demonstrating no significant skewing within the dataset. **(E)** Reanalysis of Fig 2I (impact of TEC ontogeny on the ability of TEC expression index to predict TAA immunogenicity) with downsampling of UMIs and cell counts to confirm no technical bias in the results that demonstrate fetal TECs are the most predictive of TAA immunogenicity. **(F)** Correlation between TAA-specific CD8⁺ T cell frequencies (measured by flow cytometry) and TEC expression metrics. Left, using pseudobulk expression; middle, using the TEC expression index from all TECs (top 0.2% expressors); right, using the fetal TEC expression index (top 0.01% expressors). Consistent with analyses based on CITE-seq frequencies (Fig 2D, Fig 2G, Fig 2H), the fetal TEC index showed the strongest association with TAA immunogenicity. ρ, Spearman correlation; SNR, signal-to-noise ratio. **(G)** Impact of TEC ontogeny on the ability of TEC expression indices to predict TAA immunogenicity, based on flow cytometry derived frequencies. As observed with CITE-seq data, fetal TEC expression most effectively distinguished immunogenic from poorly immunogenic TAAs. **(H)** TCRβ CDR3 length distributions for TAA-specific CD8⁺ T cells versus unsorted CD8⁺ T cells from the same donors. Individual distributions per TAA specificity shown, with dotted line representing the mean length for unsorted cells.

**Extended Data Fig 2:**
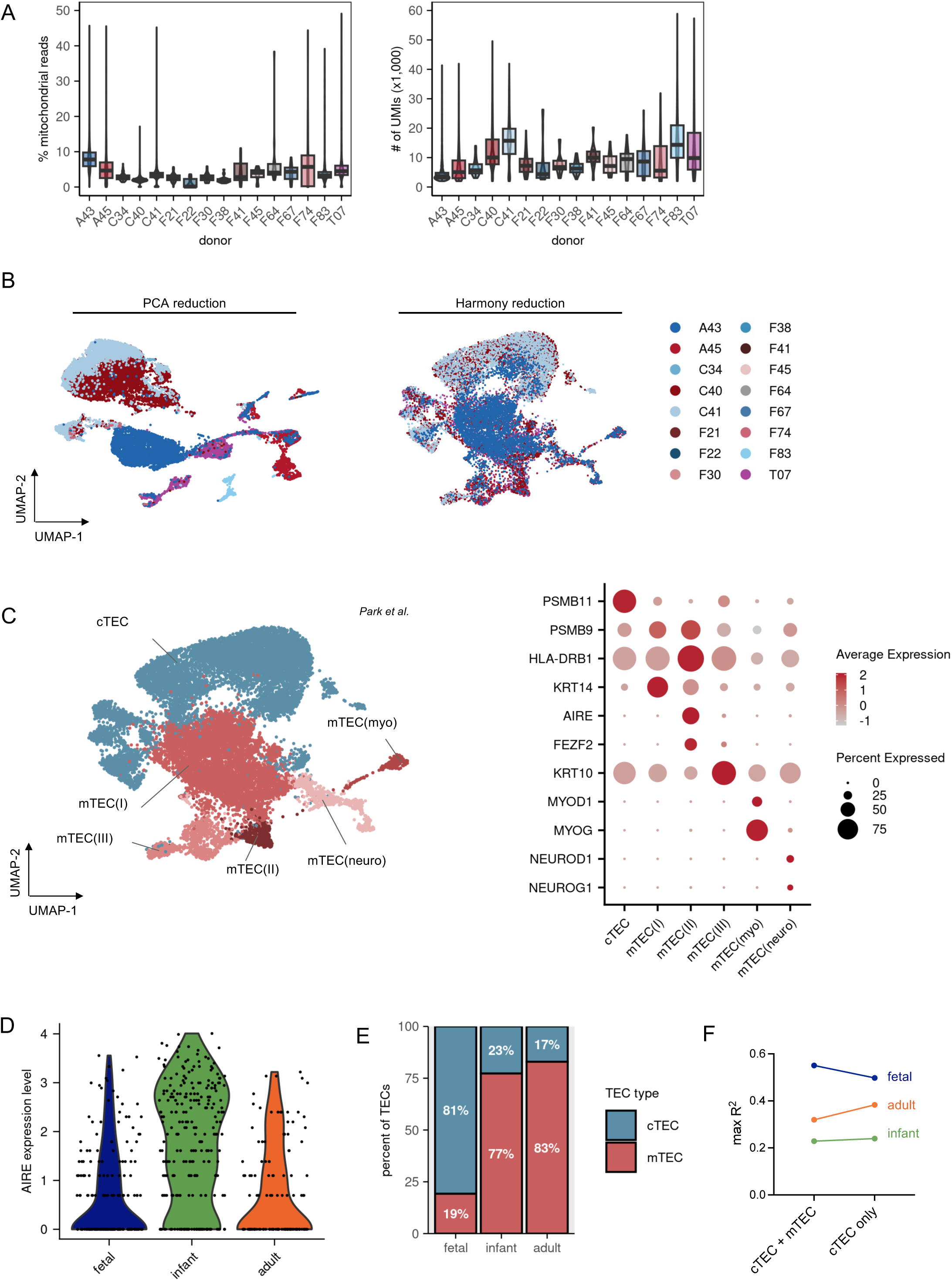
Quality control, batch correction, and characterization of thymic epithelial cell subsets across development. **(A)** Percent mitochondrial gene reads and UMIs per cell for TECs in scRNA-seq thymus dataset. **(B)** UMAP comparing PCA reduction and Harmony reduction, demonstrating removal of sample batch effect. **(C)** Assignment of TEC subsets. Left, UMAP demonstrating finalized assignments; right, dot plot of canonical genes differentiating cTECs and mTECs. **(D)** AIRE expression levels across developmental states within mTEC(II) cells, demonstrating AIRE is lowest in the fetal thymus, upregulated postnatally, and then decreases with age. **(E)** Distribution of cTEC vs mTECs by developmental state. TECs skew towards cTECs during fetal development, while the postnatal thymus has a predominance of mTECs. **(F)** Changes in maximum predictive value of TEC expression index when narrowed to only cTECs. Minimal change is noted, supporting the hypothesis that fetal cTECs may play a unique role in thymic tolerance compared to postnatal cTECs.

**Extended Data Fig 3:**
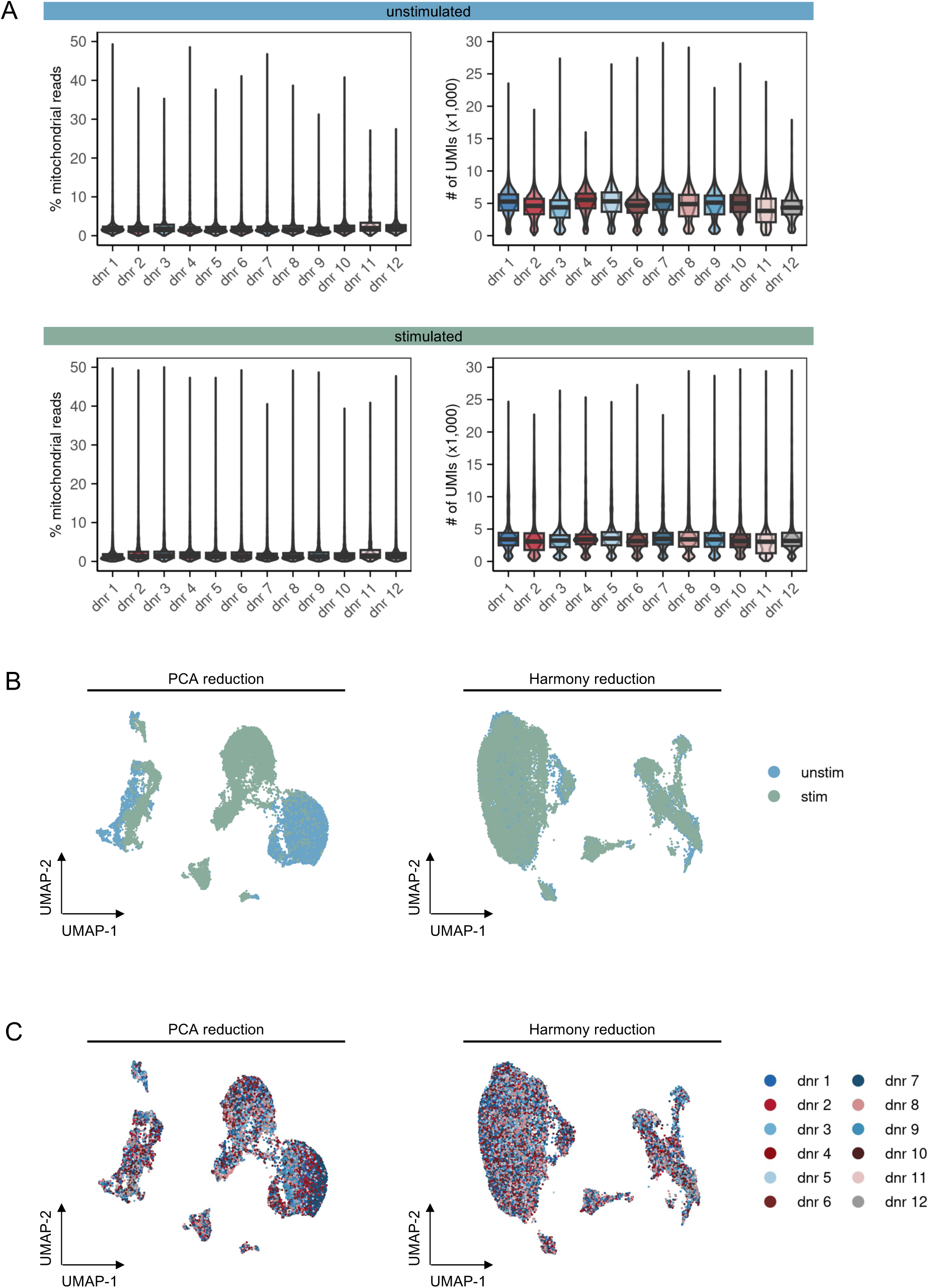
Quality control and batch correction of multimodal single cell profiling of TAA-specific CD8^+^ T cells. **(A)** Percent mitochondrial gene reads and UMIs per cell for TAA-specific CD8^+^ T cells from unstimulated, top, and stimulated, bottom, conditions. **(B)** UMAP comparison of PCA and Harmony reductions, demonstrating integration that accounts for stimulation-related effects. **(C)** UMAP comparing PCA reduction and Harmony reduction, demonstrating removal of donor batch effect.

**Extended Data Fig 4:**
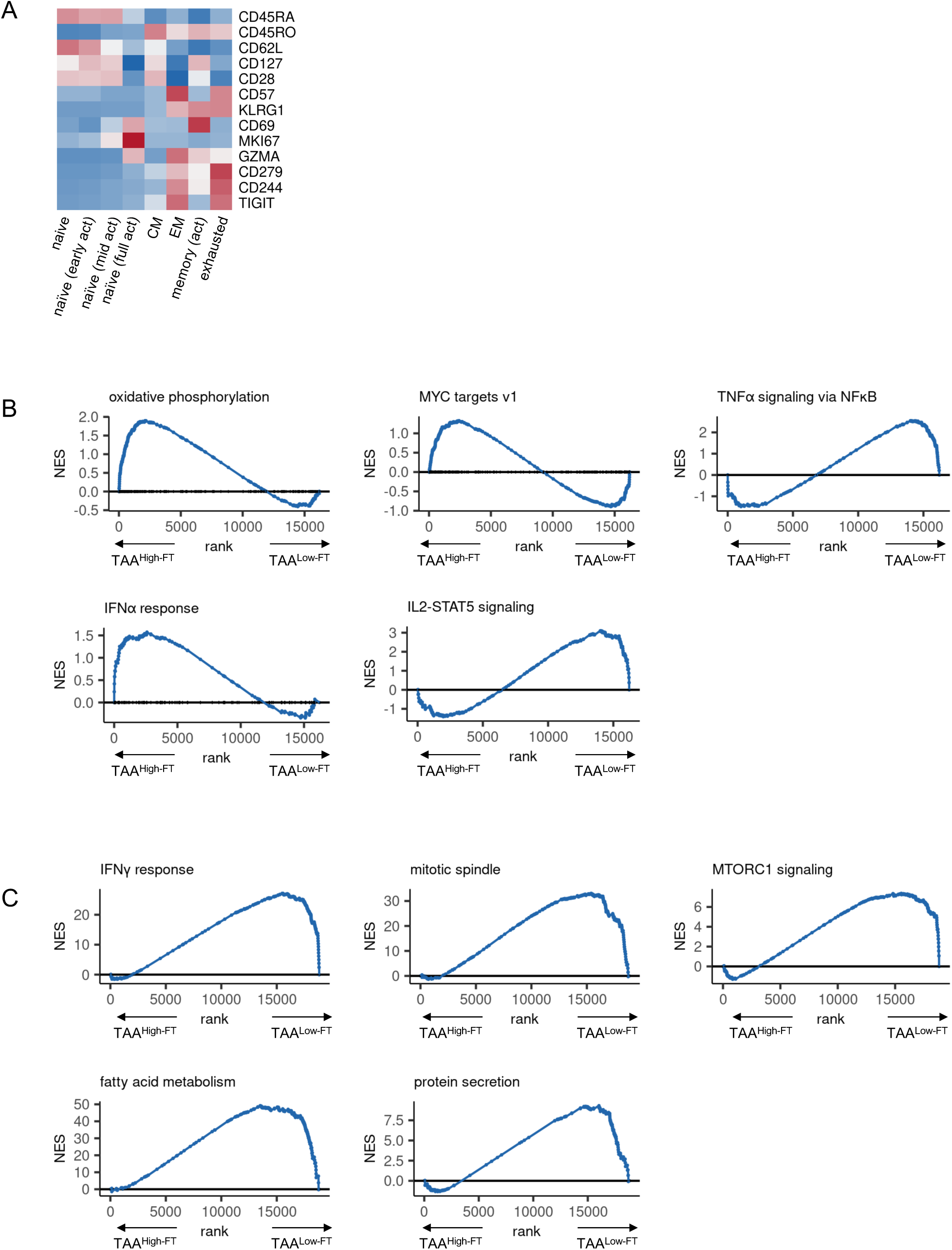
Characterization of TAA-specific CD8⁺ T cell memory states and transcriptional pathways associated with fetal TEC expression. **(A)** Expression of canonical memory markers for assignment of TAA-specific CD8^+^ T cell memory state. All markers are protein level measurements except for GZMA and MKI67, which represent RNA level measurements. **(B)** GSEA enrichment plots for those pathways found to be differentially upregulated in unstimulated CD8^+^ T cells targeting TAA^High-FT^ vs TAA^Low-FT^. **(C)** GSEA enrichment plots for those pathways found to be differentially upregulated in stimulated CD8^+^ T cells targeting TAA^High-FT^ vs TAA^Low-FT^.

**Extended Data Fig 5:**
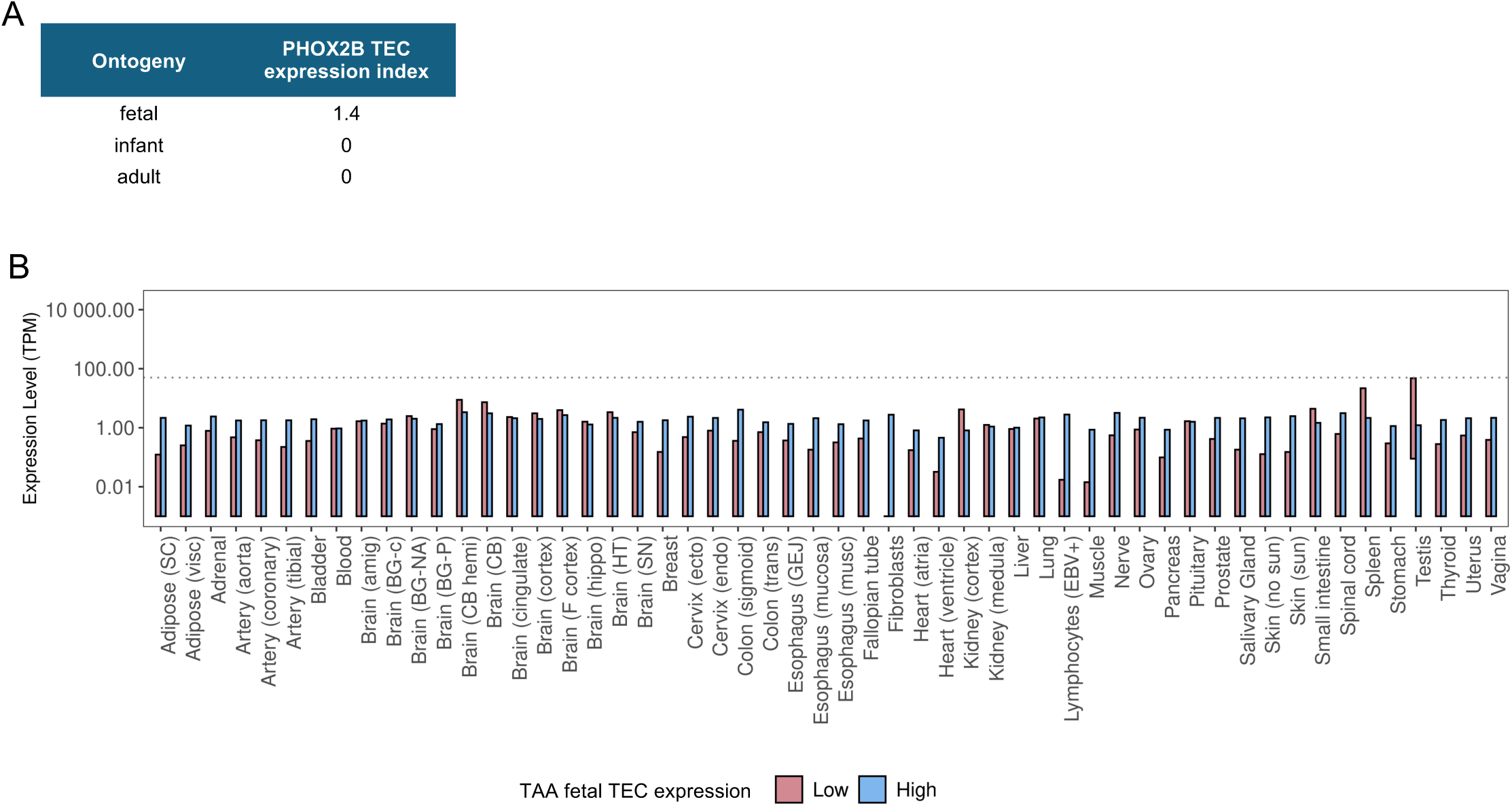
TEC expression indices for neuroblastoma candidate TAAs and their peripheral tissue distribution. **(A)** TEC expression indices for PHOX2B, previously identified candidate TAA target for neuroblastoma. PHOX2B is minimally expressed in the postnatal thymus, but highly expressed in the fetal thymus. **(B)** Peripheral tissue expression levels for neuroblastoma candidate TAAs tested, stratified by their fetal TEC expression.

## SUPPLEMENTAL INFORMATION

Supplemental Table 1. Oligotags for tetramer and donor demultiplexing of multimodal single-cell data

Supplemental Table 2. Detailed information for public melanoma RNAseq datasets

Supplemental Table 3. Sequence information for MAGE-B2 TCRs

Supplemental Fig 1. Example flow cytometry gating strategy

**Supplemental Table 1.**
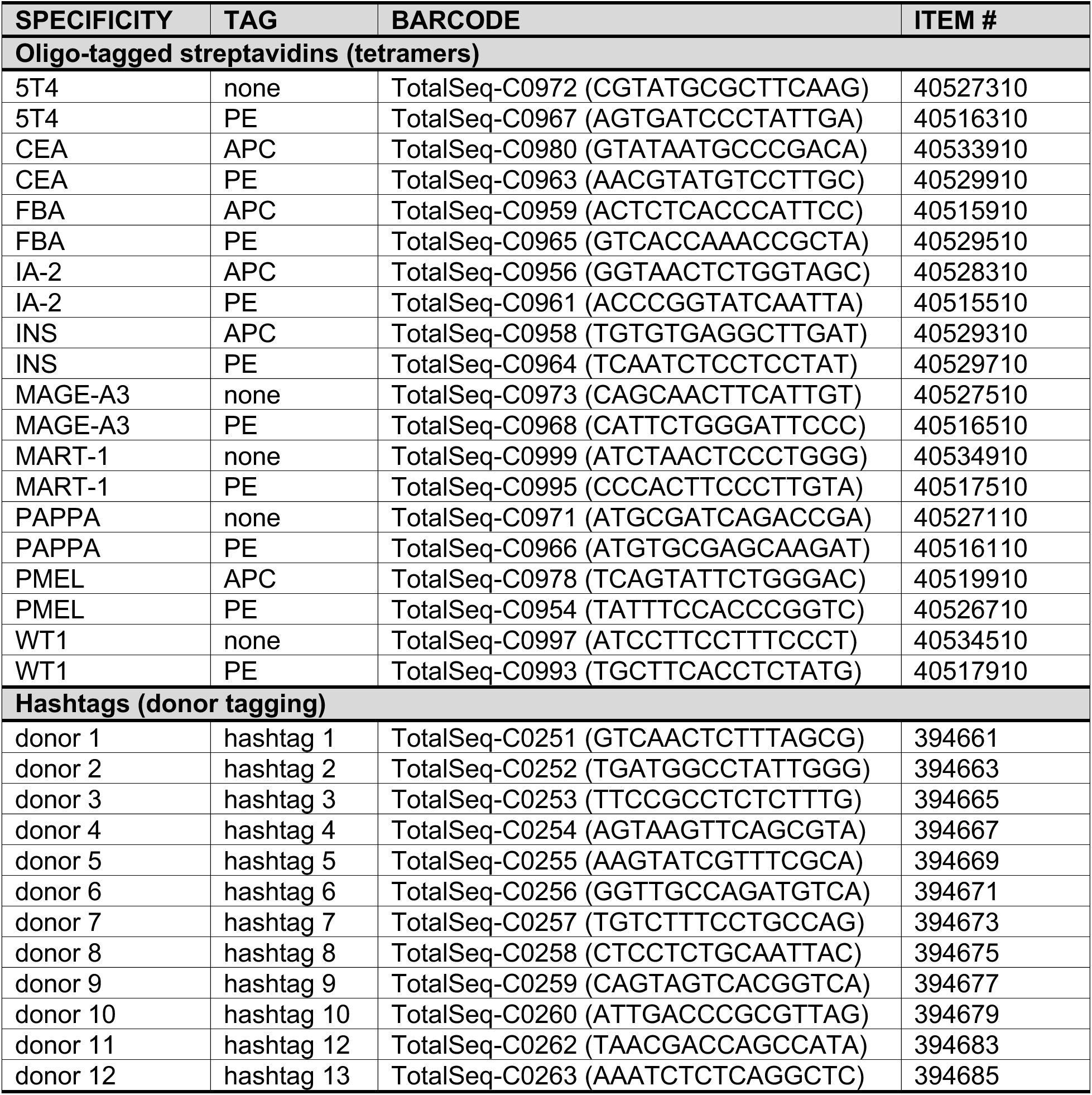
Oligotags for tetramer and donor demultiplexing of multimodal single-cell data.

**Supplemental Table 2.**
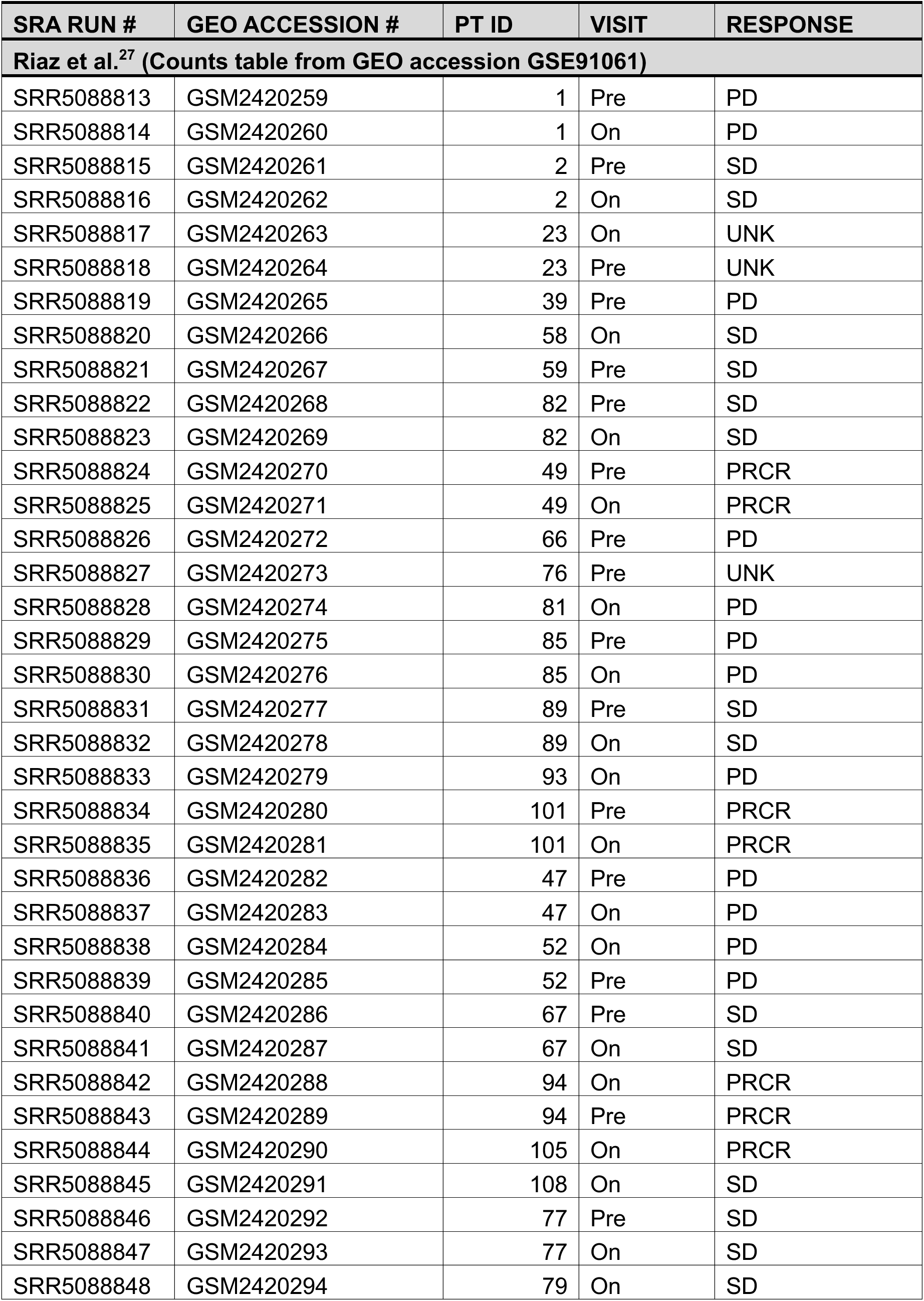
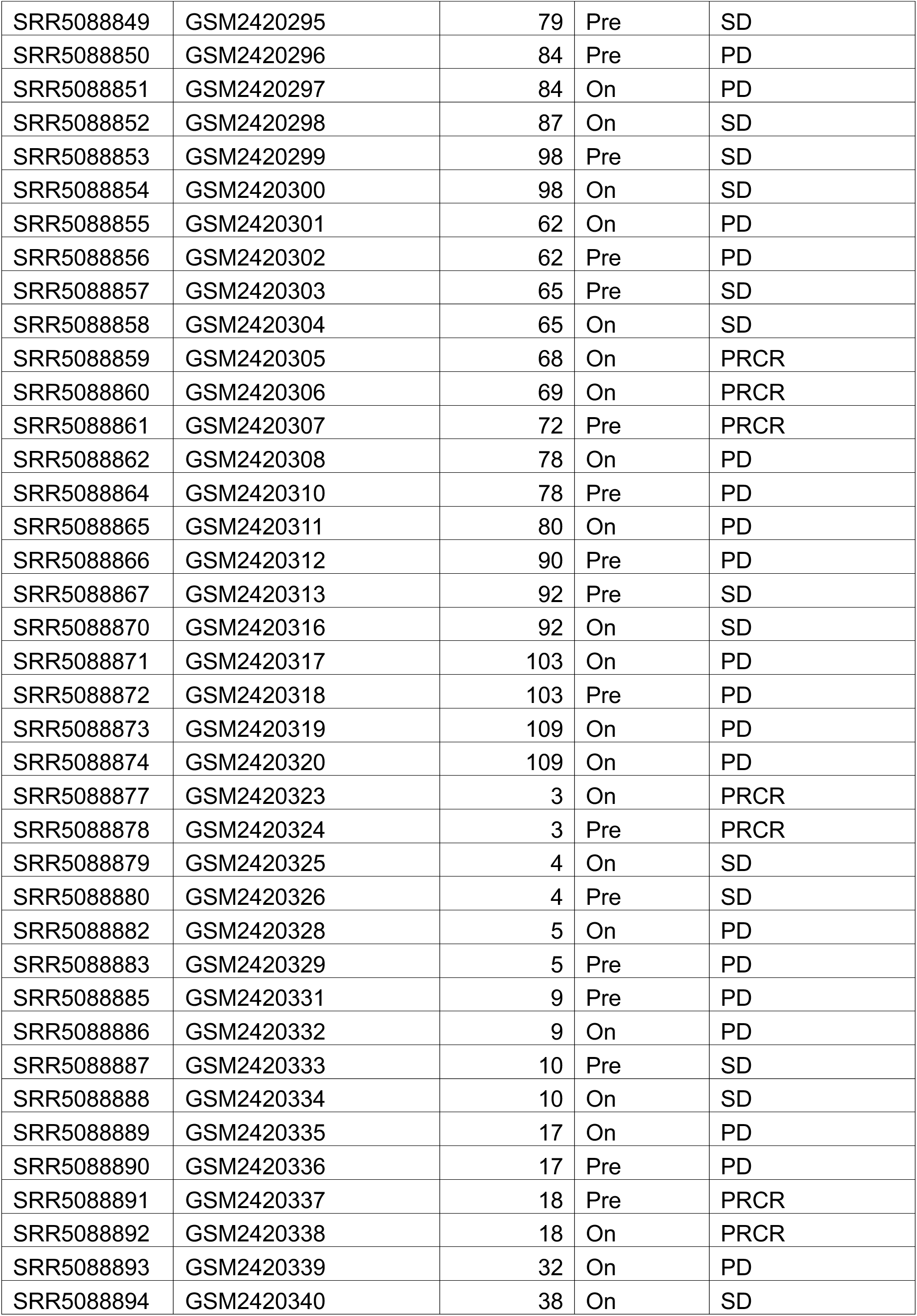
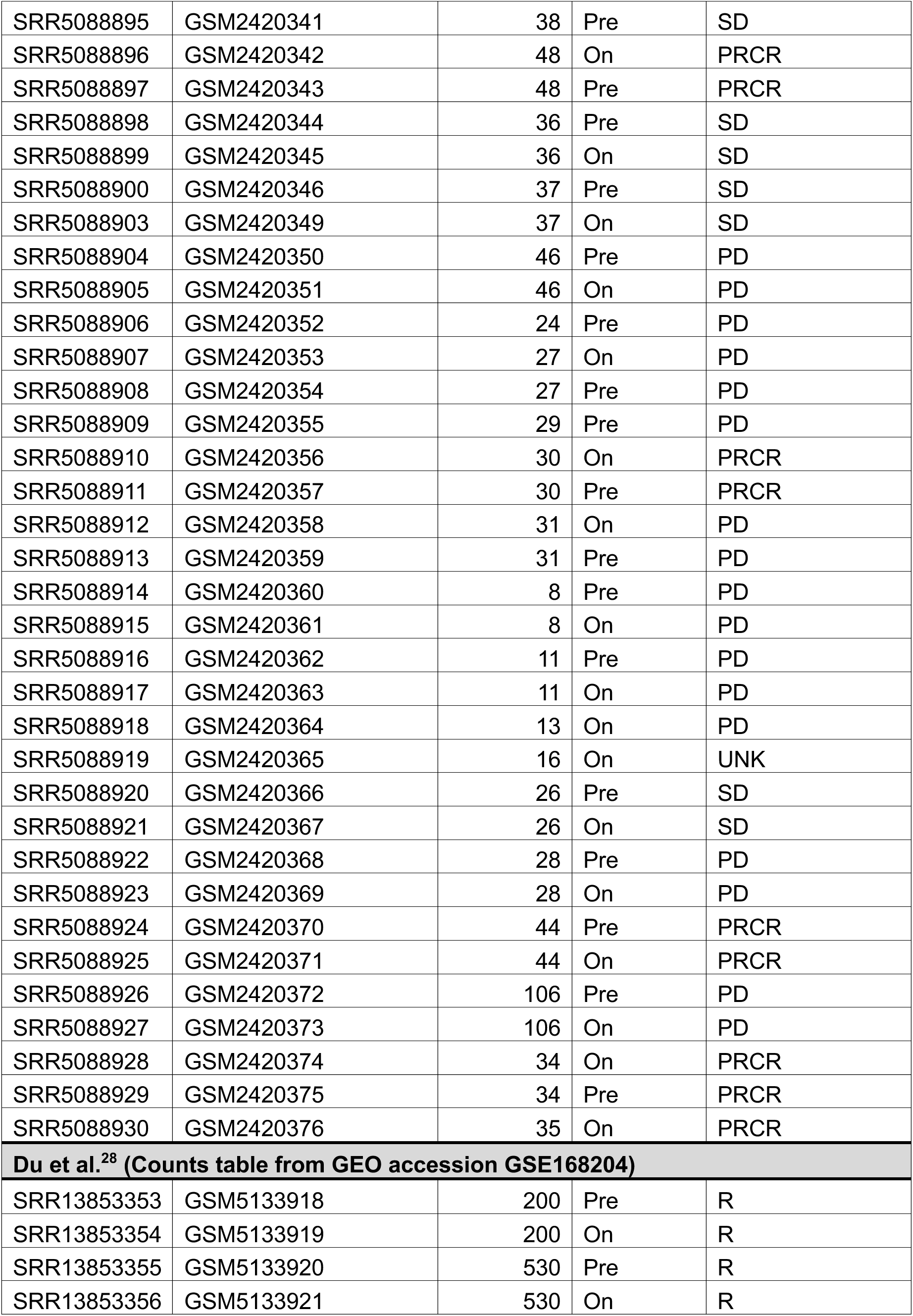
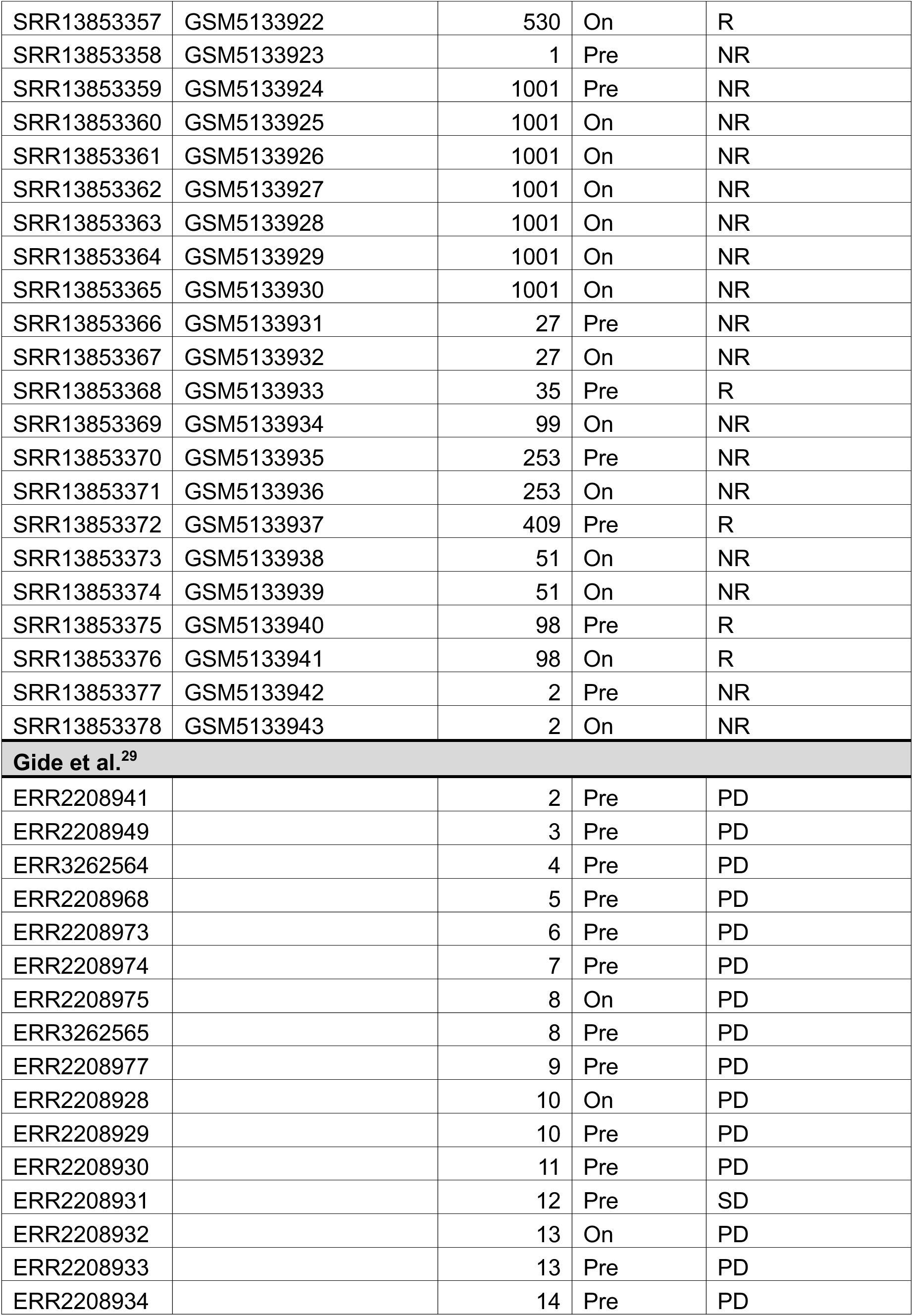
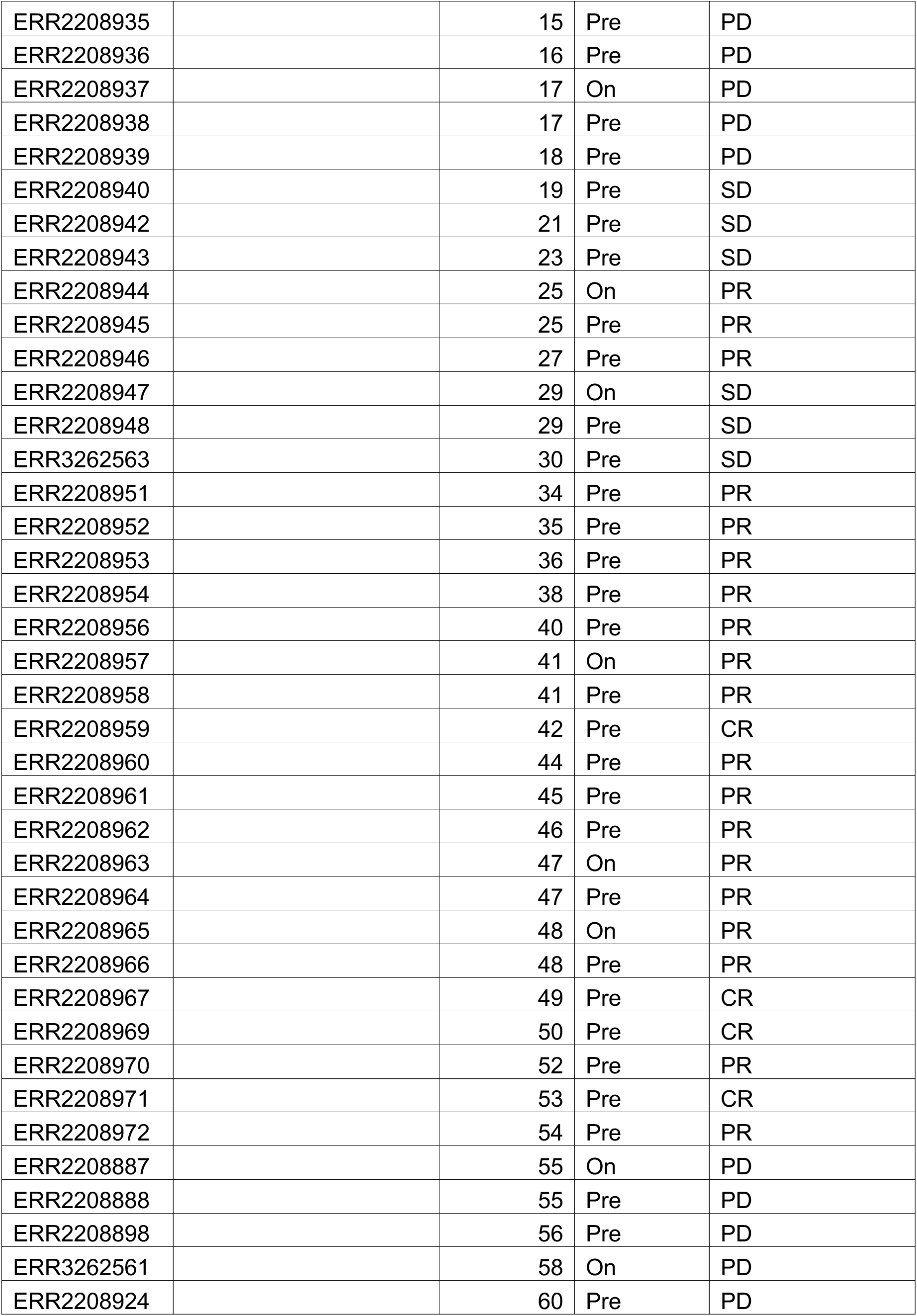
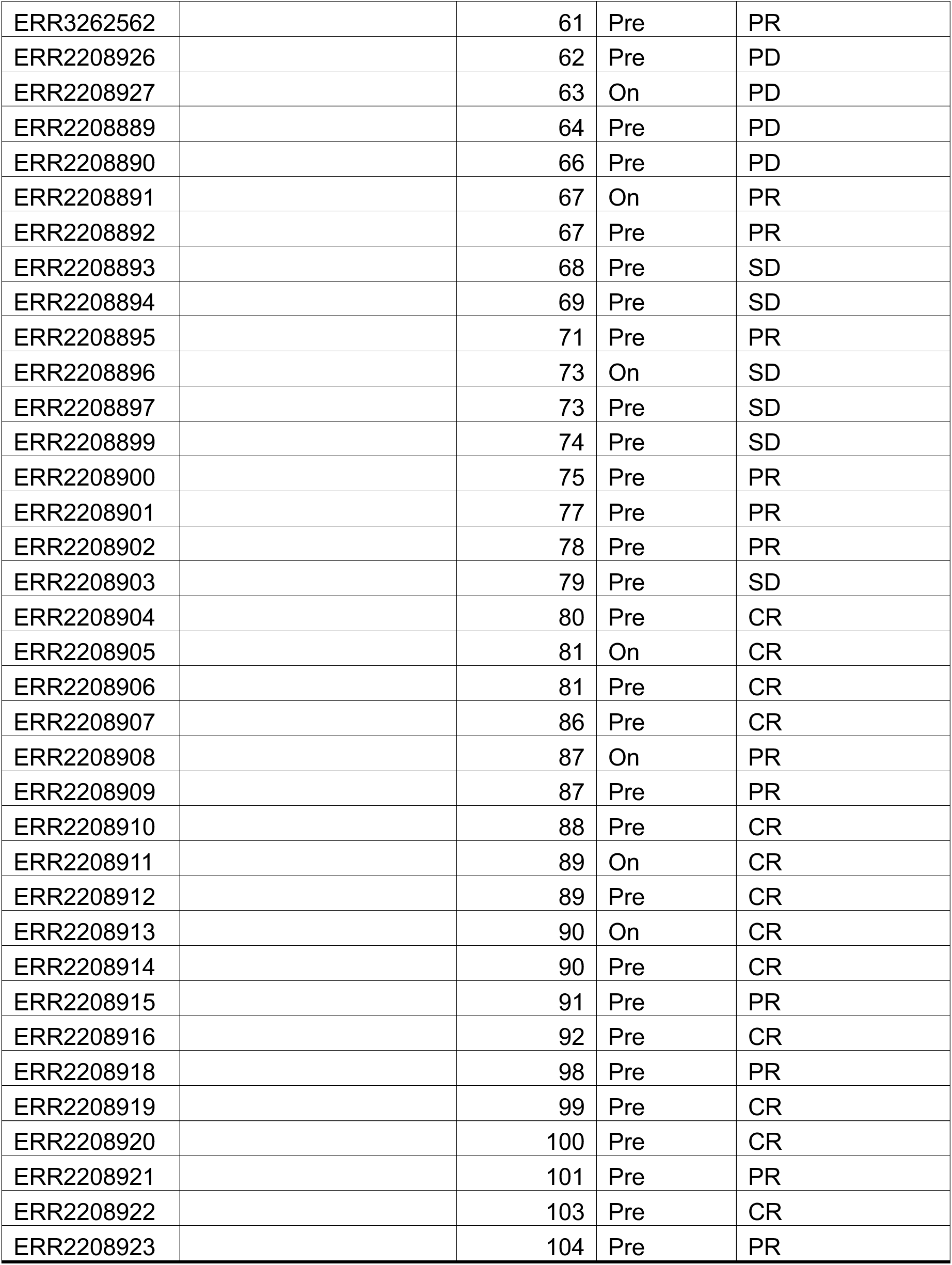
Detailed information for public melanoma RNAseq datasets.

| SRA RUN # | GEO ACCESSION # | PT ID | VISIT | RESPONSE |
| --- | --- | --- | --- | --- |
| **Riaz et al.<sup>27</sup> (Counts table from GEO accession GSE91061)** |  |  |  |  |
| SRR5088813 | GSM2420259 | 1 | Pre | PD |
| SRR5088814 | GSM2420260 | 1 | On | PD |
| SRR5088815 | GSM2420261 | 2 | Pre | SD |
| SRR5088816 | GSM2420262 | 2 | On | SD |
| SRR5088817 | GSM2420263 | 23 | On | UNK |
| SRR5088818 | GSM2420264 | 23 | Pre | UNK |
| SRR5088819 | GSM2420265 | 39 | Pre | PD |
| SRR5088820 | GSM2420266 | 58 | On | SD |
| SRR5088821 | GSM2420267 | 59 | Pre | SD |
| SRR5088822 | GSM2420268 | 82 | Pre | SD |
| SRR5088823 | GSM2420269 | 82 | On | SD |
| SRR5088824 | GSM2420270 | 49 | Pre | PRCR |
| SRR5088825 | GSM2420271 | 49 | On | PRCR |
| SRR5088826 | GSM2420272 | 66 | Pre | PD |
| SRR5088827 | GSM2420273 | 76 | Pre | UNK |
| SRR5088828 | GSM2420274 | 81 | On | PD |
| SRR5088829 | GSM2420275 | 85 | Pre | PD |
| SRR5088830 | GSM2420276 | 85 | On | PD |
| SRR5088831 | GSM2420277 | 89 | Pre | SD |
| SRR5088832 | GSM2420278 | 89 | On | SD |
| SRR5088833 | GSM2420279 | 93 | On | PD |
| SRR5088834 | GSM2420280 | 101 | Pre | PRCR |
| SRR5088835 | GSM2420281 | 101 | On | PRCR |
| SRR5088836 | GSM2420282 | 47 | Pre | PD |
| SRR5088837 | GSM2420283 | 47 | On | PD |
| SRR5088838 | GSM2420284 | 52 | On | PD |
| SRR5088839 | GSM2420285 | 52 | Pre | PD |
| SRR5088840 | GSM2420286 | 67 | Pre | SD |
| SRR5088841 | GSM2420287 | 67 | On | SD |
| SRR5088842 | GSM2420288 | 94 | On | PRCR |
| SRR5088843 | GSM2420289 | 94 | Pre | PRCR |
| SRR5088844 | GSM2420290 | 105 | On | PRCR |
| SRR5088845 | GSM2420291 | 108 | On | SD |
| SRR5088846 | GSM2420292 | 77 | Pre | SD |
| SRR5088847 | GSM2420293 | 77 | On | SD |
| SRR5088848 | GSM2420294 | 79 | On | SD |
| SRR5088849 | GSM2420295 | 79 | Pre | SD |
| SRR5088850 | GSM2420296 | 84 | Pre | PD |
| SRR5088851 | GSM2420297 | 84 | On | PD |
| SRR5088852 | GSM2420298 | 87 | On | SD |
| SRR5088853 | GSM2420299 | 98 | Pre | SD |
| SRR5088854 | GSM2420300 | 98 | On | SD |
| SRR5088855 | GSM2420301 | 62 | On | PD |
| SRR5088856 | GSM2420302 | 62 | Pre | PD |
| SRR5088857 | GSM2420303 | 65 | Pre | SD |
| SRR5088858 | GSM2420304 | 65 | On | SD |
| SRR5088859 | GSM2420305 | 68 | On | PRCR |
| SRR5088860 | GSM2420306 | 69 | On | PRCR |
| SRR5088861 | GSM2420307 | 72 | Pre | PRCR |
| SRR5088862 | GSM2420308 | 78 | On | PD |
| SRR5088864 | GSM2420310 | 78 | Pre | PD |
| SRR5088865 | GSM2420311 | 80 | On | PD |
| SRR5088866 | GSM2420312 | 90 | Pre | PD |
| SRR5088867 | GSM2420313 | 92 | Pre | SD |
| SRR5088870 | GSM2420316 | 92 | On | SD |
| SRR5088871 | GSM2420317 | 103 | On | PD |
| SRR5088872 | GSM2420318 | 103 | Pre | PD |
| SRR5088873 | GSM2420319 | 109 | On | PD |
| SRR5088874 | GSM2420320 | 109 | On | PD |
| SRR5088877 | GSM2420323 | 3 | On | PRCR |
| SRR5088878 | GSM2420324 | 3 | Pre | PRCR |
| SRR5088879 | GSM2420325 | 4 | On | SD |
| SRR5088880 | GSM2420326 | 4 | Pre | SD |
| SRR5088882 | GSM2420328 | 5 | On | PD |
| SRR5088883 | GSM2420329 | 5 | Pre | PD |
| SRR5088885 | GSM2420331 | 9 | Pre | PD |
| SRR5088886 | GSM2420332 | 9 | On | PD |
| SRR5088887 | GSM2420333 | 10 | Pre | SD |
| SRR5088888 | GSM2420334 | 10 | On | SD |
| SRR5088889 | GSM2420335 | 17 | On | PD |
| SRR5088890 | GSM2420336 | 17 | Pre | PD |
| SRR5088891 | GSM2420337 | 18 | Pre | PRCR |
| SRR5088892 | GSM2420338 | 18 | On | PRCR |
| SRR5088893 | GSM2420339 | 32 | On | PD |
| SRR5088894 | GSM2420340 | 38 | On | SD |
| SRR5088895 | GSM2420341 | 38 | Pre | SD |
| SRR5088896 | GSM2420342 | 48 | On | PRCR |
| SRR5088897 | GSM2420343 | 48 | Pre | PRCR |
| SRR5088898 | GSM2420344 | 36 | Pre | SD |
| SRR5088899 | GSM2420345 | 36 | On | SD |
| SRR5088900 | GSM2420346 | 37 | Pre | SD |
| SRR5088903 | GSM2420349 | 37 | On | SD |
| SRR5088904 | GSM2420350 | 46 | Pre | PD |
| SRR5088905 | GSM2420351 | 46 | On | PD |
| SRR5088906 | GSM2420352 | 24 | Pre | PD |
| SRR5088907 | GSM2420353 | 27 | On | PD |
| SRR5088908 | GSM2420354 | 27 | Pre | PD |
| SRR5088909 | GSM2420355 | 29 | Pre | PD |
| SRR5088910 | GSM2420356 | 30 | On | PRCR |
| SRR5088911 | GSM2420357 | 30 | Pre | PRCR |
| SRR5088912 | GSM2420358 | 31 | On | PD |
| SRR5088913 | GSM2420359 | 31 | Pre | PD |
| SRR5088914 | GSM2420360 | 8 | Pre | PD |
| SRR5088915 | GSM2420361 | 8 | On | PD |
| SRR5088916 | GSM2420362 | 11 | Pre | PD |
| SRR5088917 | GSM2420363 | 11 | On | PD |
| SRR5088918 | GSM2420364 | 13 | On | PD |
| SRR5088919 | GSM2420365 | 16 | On | UNK |
| SRR5088920 | GSM2420366 | 26 | Pre | SD |
| SRR5088921 | GSM2420367 | 26 | On | SD |
| SRR5088922 | GSM2420368 | 28 | Pre | PD |
| SRR5088923 | GSM2420369 | 28 | On | PD |
| SRR5088924 | GSM2420370 | 44 | Pre | PRCR |
| SRR5088925 | GSM2420371 | 44 | On | PRCR |
| SRR5088926 | GSM2420372 | 106 | Pre | PD |
| SRR5088927 | GSM2420373 | 106 | On | PD |
| SRR5088928 | GSM2420374 | 34 | On | PRCR |
| SRR5088929 | GSM2420375 | 34 | Pre | PRCR |
| SRR5088930 | GSM2420376 | 35 | On | PRCR |
| **Du et al.<sup>28</sup> (Counts table from GEO accession GSE168204)** |  |  |  |  |
| SRR13853353 | GSM5133918 | 200 | Pre | R |
| SRR13853354 | GSM5133919 | 200 | On | R |
| SRR13853355 | GSM5133920 | 530 | Pre | R |
| SRR13853356 | GSM5133921 | 530 | On | R |
| SRR13853357 | GSM5133922 | 530 | On | R |
| SRR13853358 | GSM5133923 | 1 | Pre | NR |
| SRR13853359 | GSM5133924 | 1001 | Pre | NR |
| SRR13853360 | GSM5133925 | 1001 | On | NR |
| SRR13853361 | GSM5133926 | 1001 | On | NR |
| SRR13853362 | GSM5133927 | 1001 | On | NR |
| SRR13853363 | GSM5133928 | 1001 | On | NR |
| SRR13853364 | GSM5133929 | 1001 | On | NR |
| SRR13853365 | GSM5133930 | 1001 | On | NR |
| SRR13853366 | GSM5133931 | 27 | Pre | NR |
| SRR13853367 | GSM5133932 | 27 | On | NR |
| SRR13853368 | GSM5133933 | 35 | Pre | R |
| SRR13853369 | GSM5133934 | 99 | On | NR |
| SRR13853370 | GSM5133935 | 253 | Pre | NR |
| SRR13853371 | GSM5133936 | 253 | On | NR |
| SRR13853372 | GSM5133937 | 409 | Pre | R |
| SRR13853373 | GSM5133938 | 51 | On | NR |
| SRR13853374 | GSM5133939 | 51 | On | NR |
| SRR13853375 | GSM5133940 | 98 | Pre | R |
| SRR13853376 | GSM5133941 | 98 | On | R |
| SRR13853377 | GSM5133942 | 2 | Pre | NR |
| SRR13853378 | GSM5133943 | 2 | On | NR |
| **Gide et al.<sup>29</sup>** |  |  |  |  |
| ERR2208941 |  | 2 | Pre | PD |
| ERR2208949 |  | 3 | Pre | PD |
| ERR3262564 |  | 4 | Pre | PD |
| ERR2208968 |  | 5 | Pre | PD |
| ERR2208973 |  | 6 | Pre | PD |
| ERR2208974 |  | 7 | Pre | PD |
| ERR2208975 |  | 8 | On | PD |
| ERR3262565 |  | 8 | Pre | PD |
| ERR2208977 |  | 9 | Pre | PD |
| ERR2208928 |  | 10 | On | PD |
| ERR2208929 |  | 10 | Pre | PD |
| ERR2208930 |  | 11 | Pre | PD |
| ERR2208931 |  | 12 | Pre | SD |
| ERR2208932 |  | 13 | On | PD |
| ERR2208933 |  | 13 | Pre | PD |
| ERR2208934 |  | 14 | Pre | PD |
| ERR2208935 |  | 15 | Pre | PD |
| ERR2208936 |  | 16 | Pre | PD |
| ERR2208937 |  | 17 | On | PD |
| ERR2208938 |  | 17 | Pre | PD |
| ERR2208939 |  | 18 | Pre | PD |
| ERR2208940 |  | 19 | Pre | SD |
| ERR2208942 |  | 21 | Pre | SD |
| ERR2208943 |  | 23 | Pre | SD |
| ERR2208944 |  | 25 | On | PR |
| ERR2208945 |  | 25 | Pre | PR |
| ERR2208946 |  | 27 | Pre | PR |
| ERR2208947 |  | 29 | On | SD |
| ERR2208948 |  | 29 | Pre | SD |
| ERR3262563 |  | 30 | Pre | SD |
| ERR2208951 |  | 34 | Pre | PR |
| ERR2208952 |  | 35 | Pre | PR |
| ERR2208953 |  | 36 | Pre | PR |
| ERR2208954 |  | 38 | Pre | PR |
| ERR2208956 |  | 40 | Pre | PR |
| ERR2208957 |  | 41 | On | PR |
| ERR2208958 |  | 41 | Pre | PR |
| ERR2208959 |  | 42 | Pre | CR |
| ERR2208960 |  | 44 | Pre | PR |
| ERR2208961 |  | 45 | Pre | PR |
| ERR2208962 |  | 46 | Pre | PR |
| ERR2208963 |  | 47 | On | PR |
| ERR2208964 |  | 47 | Pre | PR |
| ERR2208965 |  | 48 | On | PR |
| ERR2208966 |  | 48 | Pre | PR |
| ERR2208967 |  | 49 | Pre | CR |
| ERR2208969 |  | 50 | Pre | CR |
| ERR2208970 |  | 52 | Pre | PR |
| ERR2208971 |  | 53 | Pre | CR |
| ERR2208972 |  | 54 | Pre | PR |
| ERR2208887 |  | 55 | On | PD |
| ERR2208888 |  | 55 | Pre | PD |
| ERR2208898 |  | 56 | Pre | PD |
| ERR3262561 |  | 58 | On | PD |
| ERR2208924 |  | 60 | Pre | PD |
| ERR3262562 |  | 61 | Pre | PR |
| ERR2208926 |  | 62 | Pre | PD |
| ERR2208927 |  | 63 | On | PD |
| ERR2208889 |  | 64 | Pre | PD |
| ERR2208890 |  | 66 | Pre | PD |
| ERR2208891 |  | 67 | On | PR |
| ERR2208892 |  | 67 | Pre | PR |
| ERR2208893 |  | 68 | Pre | SD |
| ERR2208894 |  | 69 | Pre | SD |
| ERR2208895 |  | 71 | Pre | PR |
| ERR2208896 |  | 73 | On | SD |
| ERR2208897 |  | 73 | Pre | SD |
| ERR2208899 |  | 74 | Pre | SD |
| ERR2208900 |  | 75 | Pre | PR |
| ERR2208901 |  | 77 | Pre | PR |
| ERR2208902 |  | 78 | Pre | PR |
| ERR2208903 |  | 79 | Pre | SD |
| ERR2208904 |  | 80 | Pre | CR |
| ERR2208905 |  | 81 | On | CR |
| ERR2208906 |  | 81 | Pre | CR |
| ERR2208907 |  | 86 | Pre | CR |
| ERR2208908 |  | 87 | On | PR |
| ERR2208909 |  | 87 | Pre | PR |
| ERR2208910 |  | 88 | Pre | CR |
| ERR2208911 |  | 89 | On | CR |
| ERR2208912 |  | 89 | Pre | CR |
| ERR2208913 |  | 90 | On | CR |
| ERR2208914 |  | 90 | Pre | CR |
| ERR2208915 |  | 91 | Pre | PR |
| ERR2208916 |  | 92 | Pre | CR |
| ERR2208918 |  | 98 | Pre | PR |
| ERR2208919 |  | 99 | Pre | CR |
| ERR2208920 |  | 100 | Pre | CR |
| ERR2208921 |  | 101 | Pre | PR |
| ERR2208922 |  | 103 | Pre | CR |
| ERR2208923 |  | 104 | Pre | PR |

**Supplemental Table 3.**
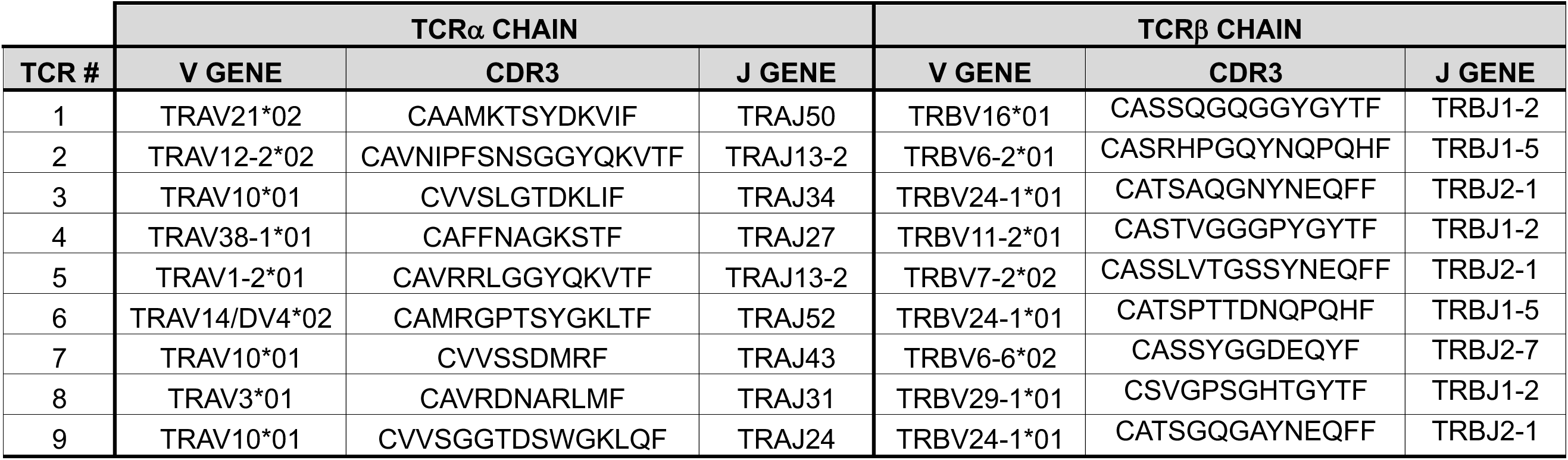
Sequence information for MAGE-B2 TCRs. ^38–39^

**Supplemental Fig 1:**
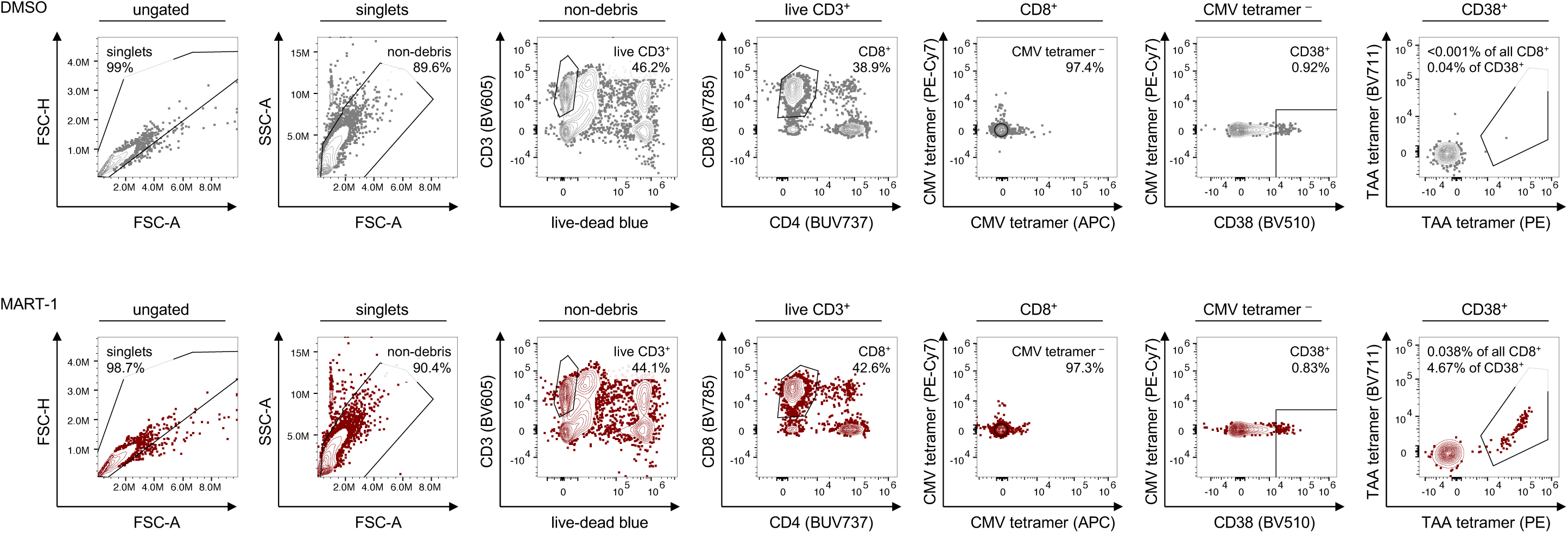
Example flow cytometry gating strategy. Healthy donor PBMCs were stimulated with TAA peptides, cultured for 7 days, and antigen-specific responses measured with peptide-MHC tetramers using flow cytometry. Activated, antigen specific CD8^+^ T cells defined by gating: singlets ➜ non-debris ➜ live CD3^+^ T cells ➜ CD8^+^ T cells ➜ CMV tetramer^−^ ➜ CD38^+^ ➜ TAA tetramer^+^. Example gating for DMSO stimulated and MART-1 stimulated cultures.

